# *Gli1*+ mesenchymal stromal cells modulate epithelial metaplasia in lung fibrosis

**DOI:** 10.1101/841957

**Authors:** Monica Cassandras, Chaoqun Wang, Jaymin Kathiriya, Tatsuya Tsukui, Peri Matatia, Michael Matthay, Paul Wolters, Ari Molofsky, Dean Sheppard, Hal Chapman, Tien Peng

## Abstract

Organ fibrosis is often accompanied by aberrant epithelial reprogramming, culminating in a transformed barrier composed of scar and metaplastic epithelium. Understanding how the scar promotes an abnormal epithelial response could better inform strategies to reverse the fibrotic damage. Here we show that *Gli1*+ mesenchymal stromal cells (MSCs), previously shown to contribute to myofibroblasts in the scar, promote metaplastic differentiation of airway progenitors into KRT5+ basal cells *in vitro* and *in vivo*. During fibrotic repair, *Gli1*+ MSCs integrate hedgehog activation to promote metaplastic KRT5 differentiation by upregulating BMP antagonism in the progenitor niche. Restoring the balance towards BMP activation attenuated metaplastic KRT5+ differentiation while promoting adaptive alveolar differentiation. Finally, fibrotic human lungs demonstrate altered BMP activation in the metaplastic epithelium. These findings show that *Gli1*+ MSCs integrate hedgehog signaling as a rheostat to control BMP activation in the progenitor niche to determine regenerative outcome in fibrosis.

**Highlights:**

- *Gli1*+ MSCs are required for metaplastic airway progenitor differentiation into KRT5+ basal cells.
- Hedgehog activation of MSCs promotes KRT5 differentiation of airway progenitors by suppressing BMP activation.
- Restoring BMP activation attenuates metaplastic KRT5 differentiation
- Metaplastic KRT5+ basal cells in human fibrotic lungs demonstrate altered BMP activation.

## Introduction

A canonical feature of wound repair is the formation of scar tissue, composed of inflammatory cells and resident mesenchymal subsets that dynamically remodel the wound to close any barrier gaps. Scars are sometimes accompanied by metaplasia of adjacent epithelium, marked by conversion of one mature cell type into another that is not typically present at the site of injury (Giroux and Rustgi, 2017). Metaplasia is present in numerous pathological contexts ranging from the relatively benign (*e.g.* Barrett’s esophagus) to the extremely morbid (*e.g.* Idiopathic pulmonary fibrosis, or IPF). One of the cardinal features of IPF, the most prevalent and deadly subtype of progressive fibrotic lung diseases, is the appearance of “bronchiolization” on histology (Chilosi et al., 2002). This pathological feature denotes the ectopic appearance of proximal bronchus/airway epithelium within the distal lung, characterized by metaplastic KRT5+ basal cells lining cystic airspaces in the alveoli alongside patches of fibroblastic scars (Seibold et al., 2013; Xu et al., 2016). While the functional relationship between metaplasia and scarring is unclear in IPF, the appearance of metaplastic KRT5+ cells is correlated with worsening disease severity and survival (Prasse et al., 2019). This suggests that epithelial metaplasia is a clinically relevant feature of organ fibrosis and a potential therapeutic target.

The lung resembles an upside-down tree with distinct compartments along the proximal-distal axis that are populated by unique resident epithelial stem/progenitor cells. KRT5+ basal cells reside in the most proximal end in the trachea and large airways to give rise to airway epithelial lineages, and SFTPC+ Type 2 cells reside in the distal alveolar sacs to generate functional alveolar epithelium (Hogan et al., 2014). In recent years, numerous studies have shown that KRT5-negative progenitors from the airway can migrate and reconstitute epithelium in the alveoli in response to severe damage (Ray et al., 2016; Vaughan et al., 2015; Yang et al., 2018; Yee et al., 2017). While the heterogeneity within this airway progenitor population (EpCAM+ cells expressing combinations of *Sox2*, *tp63*, *Scgb1a1*, and *Itgb4*) has not been fully resolved, genetic lineage tracing and transplant experiments clearly demonstrate that KRT5-negative airway progenitors can replete the damaged alveolar surface with endogenous (SFTPC+) or metaplastic (KRT5+) epithelium. This dichotomous path to adaptive or metaplastic differentiation has significant functional consequences, but it is unclear how cell-extrinsic factors in the stem/progenitor cell niche modulate the regenerative outcome.

*Gli1*, a transcriptional readout of hedgehog (Hh) activation, labels mesenchymal cells adjacent to the airway epithelium (Peng et al., 2015). *Gli1*+ cells have been demonstrated to exhibit properties of mesenchymal stromal cells (MSCs) *ex vivo (Kramann et al., 2015; Zhao et al., 2015; Zhao et al., 2014)*, and contribute to fibrotic scarring in the lung and other organs through differentiation into myofibroblasts and secretion of collagen (Kramann et al., 2015). Despite the close proximity of *Gli1*+ mesenchymal cells to airway progenitors, their functional interactions remain undefined. We have previously demonstrated that ectopic activation of hedgehog in the distal alveolar mesenchyme disrupts SFTPC+ alveolar progenitor renewal during homeostasis (Wang et al., 2018). This suggests that Hh activation of *Gli1*+ mesenchymal cells can alter epithelial progenitor fate by modifying niche factors during fibrotic repair. In this study, we set out to define the functional interactions between the fibrotic scar and metaplastic differentiation of epithelial progenitors in the fibrotic lung.

## Results

### *Gli1*+ mesenchymal cells promote metaplastic KRT5 differentiation from the airway

SOX2+ progenitors are normally restricted to the airway epithelium during homeostasis, but they migrate into the alveoli during severe alveolar damage from influenza (Ray et al., 2016; Xi et al., 2017). To determine whether SOX2+ progenitors have similar migratory capacity in a fibrotic model, we adapted a repetitive bleomycin model of lung fibrosis that better captured features of IPF compared to the single dose model, including the appearance of cystic airspaces in the alveoli (Degryse et al., 2010; Kurche et al., 2019). Our genetic lineage tracing confirms that *Sox2* lineage traced (*Sox2* Lin+) airway progenitors expand and differentiate into metaplastic KRT5+/SOX2+ or endogenous SFTPC+/SOX2-epithelium in the alveoli in this model of severe fibrotic injury (Figure 1A model, Figure S1A). The ectopic appearance of KRT5+ cells includes both in the alveoli where KRT5+ cells form cystic structures within the alveoli, as well as in the distal airways close to the bronchoalveolar ductal junction (BADJ) where KRT5+ cells are normally absent. KRT5+ and SFTPC+ cells form spatially-segregated patches within the damaged alveoli after fibrotic injury, with the airway SOX2 marker co-localizing with KRT5 but not SFTPC (Figure 1B).

**Figure 1.**
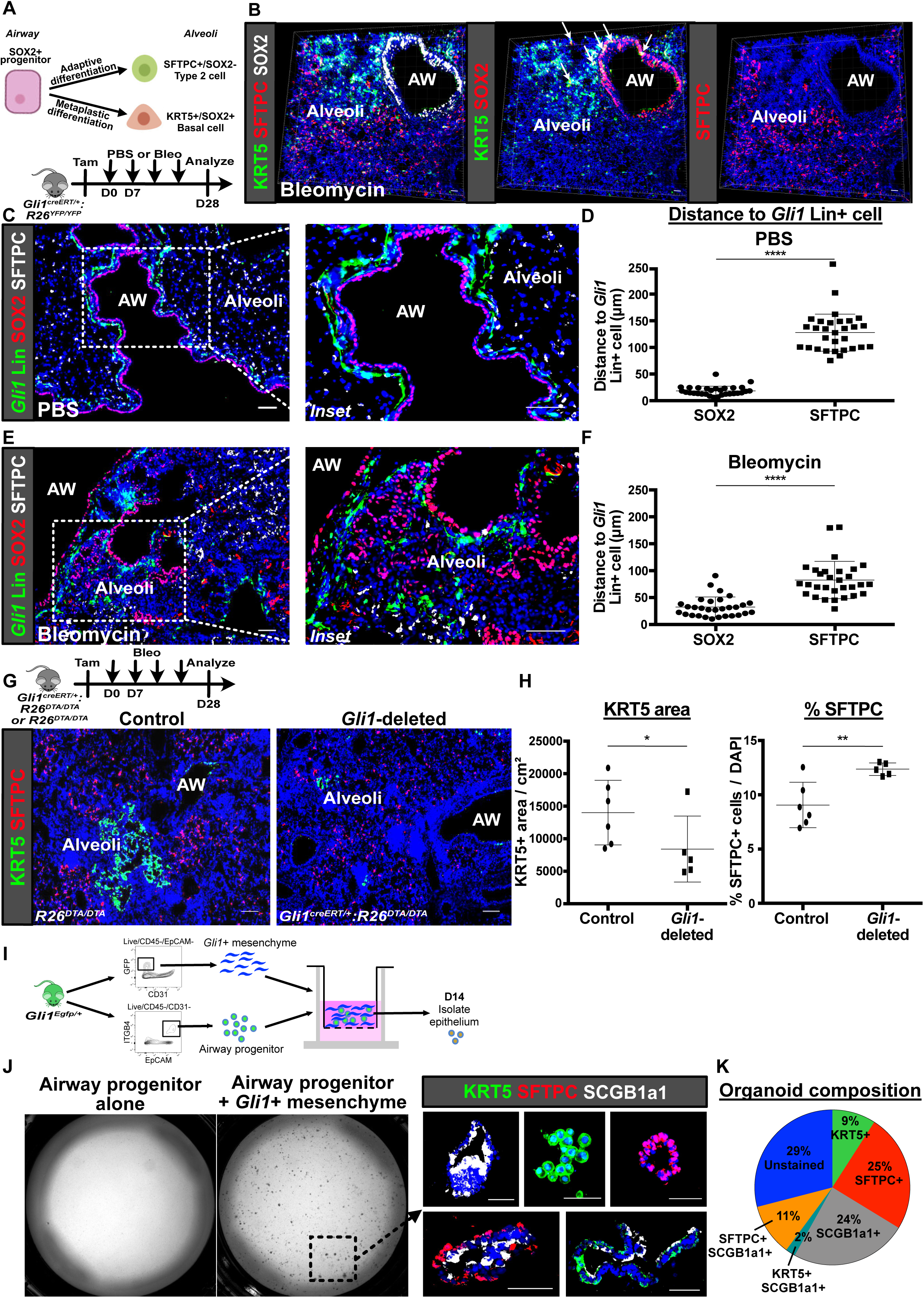
*Gli1*+ mesenchymal cells promote metaplastic KRT5 differentiation from the airway. (A) Current model of airway progenitors migrating into the alveoli to form endogenous SFTPC+ Type 2 cells or metaplastic KRT5+ basal cells after severe injury. (B) 3D thick slice imaging shows overlap of SOX2 and KRT5 in the metaplastic basal cells in the alveoli after injury, and lack of SOX2 expression in the SFTPC+ cells. (C) Histology of proximally-located *Gli1*-lineage traced (*Gli1* Lin+) mesenchymal cells and SOX2+ airway progenitors during homeostasis (PBS treated). (D) Average cell-to-cell distance (individual distances averaged per field) shows proximity of SOX2+ progenitors and SFTPC+ alveolar progenitors respectively to *Gli1* Lin+ cells during homeostasis. Data are expressed as mean ± SD. (E) Histology shows migration of *Gli1* Lin+ mesenchymal cells into the alveoli close to alveolar SOX2+ progenitors during fibrotic repair (Bleomycin treated). (F) Average cell-to-cell distance shows proximity of SOX2+ progenitors and SFTPC+ alveolar progenitors respectively to *Gli1* Lin+ cells during fibrotic repair. Data are expressed as mean ± SD. (G) Histology of the lung shows extent of metaplastic KRT5 differentiation after genetic deletion of *Gli1*+ cells followed by fibrotic repair. (H) Histology quantification reveals genetic deletion of *Gli1*+ cells resulted in smaller areas of the lung containing KRT5+ pods and larger percentage of SFTPC+ cells in the alveoli with bleomycin injury. Data are expressed as mean ± SD. (I) Model of 3D organoid assay of airway progenitor differentiation by co-culturing sorted *Gli1*+ cells and ITGB4+ airway progenitors in matrigel on an air-liquid interface. (J) Airway progenitors cultured alone fail to form organoids, while airway progenitors co-cultured with *Gli1*+ mesenchyme form organoids that demonstrate multi-lineage specification. (K) Cellular composition of individual airway progenitor-derived organoids. AW = airway. Scale bars, 100 μm. See also Figure S1 and S2.

We have previously demonstrated that the airway epithelium secretes sonic hedgehog (SHH), which promotes the expression of *Gli1* in the mesenchymal cells adjacent to the airway progenitors (Peng et al., 2015). Here, immunohistochemistry of *Gli1* lineage-labeled cells (*Gli1* Lin+) demonstrates that they are localized right beneath the basement membrane in close proximity to SOX2+ airway epithelium (Figure 1C,D) during normal homeostasis. Similar to the SOX2+ progenitors, *Gli1* Lin+ mesenchymal cells also respond to fibrotic injury by migrating into the alveolar compartment (Figure 1E, Figure S1B). Cell-to-cell distance quantification shows that the ectopic SOX2+ cells in the alveoli maintain a close distance to *Gli1* Lin+ cells that migrated into the alveoli (Figure 1F), suggesting a functional interaction between migratory *Gli1* Lin+ mesenchyme and SOX2+ progenitors during alveolar repair.

To determine whether *Gli1*+ mesenchymal cells are required for metaplastic KRT5 differentiation in the alveoli, we genetically deleted *Gli1*+ cells by inducible expression of diphtheria toxin A (DTA) within *Gli1*-expressing cells. Tamoxifen induction of *Gli1^creERT/+:^R26^DTA/DTA^* (*Gli1*-deleted) animals followed by repetitive bleomycin resulted in significantly reduced areas of metaplastic KRT5+ pods on histology compared to control *R26^DTA/DTA^* animals (Figure 1G,H). Conversely, a significantly higher proportion of the alveolar epithelium is marked by SFTPC, a Type 2 cell marker, in the *Gli1*-deleted lungs (Figure 1G,H). This suggests that activated *Gli1*+ mesenchymal cells mobilize and promote metaplastic differentiation of airway progenitors into KRT5+ lineage in the damaged alveoli, at the expense of endogenous differentiation into SFTPC+ cells.

To determine the ability of *Gli1*+ mesenchymal cells to modify airway progenitors *in vitro*, we co-cultured isolated airway progenitors with or without *Gli1*+ mesenchymal cells (isolated from uninjured lungs of *Gli1^EGFP^* reporter) in a heterotypic 3D organoid assay with minimal supplement on an air-liquid interface (Wang et al., 2018)(Figure 1I). ITGB4 is a cell surface marker of airway progenitors that largely overlaps with SOX2 (Figure S2A), and utilized to sort for airway progenitors in transplantation experiments where they give rise to both KRT5+ and SFTPC+ epithelium in damaged lungs (Vaughan et al., 2015). ITGB4+ progenitors cultured without mesenchyme failed to form organoids in the minimal-supplemented condition, but surprisingly, co-culture of ITGB4+ airway progenitors with *Gli1*+ mesenchyme resulted in significant growth in the 3D matrigel (Figure 1J). Histologic analysis of the organoids co-cultured with *Gli1*+ cells demonstrated heterogeneous lineage differentiation that included organoids containing either KRT5+ or SFTPC+ cells, but never both in the same organoid. The plurality of the organoids contained SCGB1A1+ cells, a Cub cell marker, including organoids that are both SCGB1A1+/SFTPC+ and SCGB1A1+/KRT5+ (Figure 1K), suggesting that KRT5+ and SFTPC+ cells can be derived from a SCGB1A1+ precursor. This demonstrates that isolated *Gli1*+ mesenchyme form a niche to support airway progenitor expansion and differentiation *in vitro*, and our heterotypic organoid model presents an attractive platform to assay putative niche factors that regulate metaplastic (KRT5) vs. adaptive (SFTPC) differentiation from airway progenitors.

### Mesenchymal Hh activation promotes metaplastic KRT5 differentiation

*Gli1*+ mesenchyme integrates extracellular cues during normal homeostasis and repair, including the availability of the SHH ligand that modulates the level of Hh activation in the tissue microenvironment (Peng et al., 2015). To determine the role of mesenchymal Hh activation in modulating airway progenitor differentiation, we inducibly expressed a constitutively active form of the Hh effector, *Smo* (*SmoM2*), in *Gli1*+ cells (Jeong et al., 2004). Over-activation of Hh in *Gli1*+ mesenchyme through tamoxifen induction of *Gli1^creERT/+:^R26^SmoM2/YFP^* (referred to as Hh-activated) animals did not result in metaplastic differentiation of KRT5+ cells in the uninjured lung (Figure S3A,B). However, when Hh over-activation is followed by fibrotic injury with bleomycin, Hh-activated animals demonstrated a significant increase in areas of the lung with metaplastic KRT5+ basal cells compared to controls (*Gli1^creERT/+:^R26^YFP/YFP^*) (Figure 2A,B), along with an increase in the ratio of *Krt5*/*SFTPC* transcripts in the whole lung (Figure S3C). Correspondingly, Hh-activated lungs contain significantly reduced areas of functional SFTPC+ alveolar epithelium (Figure 2A,B). This data suggests that Hh activation within *Gli1*+ mesenchyme, in conjunction with a fibrotic stimulus, acts in trans to promote the metaplastic differentiation of KRT5+ cells from airway progenitors.

**Figure 2.**
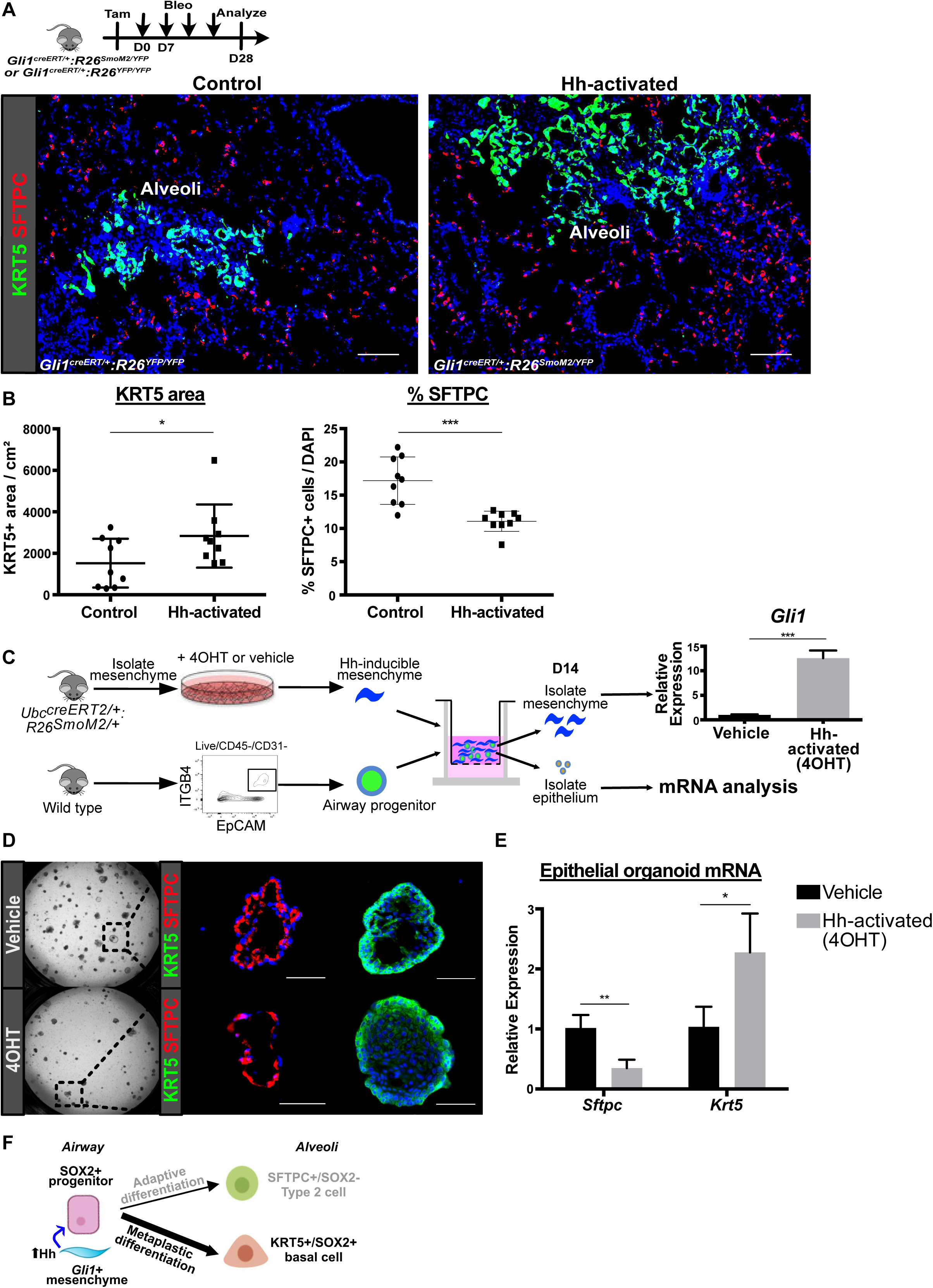
Mesenchymal Hh activation promotes metaplastic KRT5 differentiation. (A) Histology shows extent of metaplastic KRT5 differentiation following constitutive activation of Hh signaling in *Gli1* Lin+ cells followed by fibrotic repair. (B) Histology quantification shows constitutive Hh activation in *Gli1*+ cells resulted in larger areas of the lung containing KRT5+ pods and smaller percentage of SFTPC+ cells in the alveoli with bleomycin injury. Data are expressed as mean ± SD. (C) Model of 3D airway organoid assay using Hh-inducible mesenchyme whereby pretreatment of 4-hydroxytamoxifen (4OHT) induces Hh activation in the mesenchyme as shown by upregulation of *Gli1* transcript compared to vehicle (ethanol). (D) Histology of airway progenitor-derived organoids co-cultured with Hh-activated mesenchyme. (E) qPCR of isolated epithelium after 14 day co-culture shows that mesenchymal Hh-activation reduces the expression of *SFTPC* while promoting *Krt5* expression in the airway progenitor-derived organoids. Data are expressed as mean ± SD. (F) Model of Hh activation in the *Gli1*+ mesenchyme acting in trans to promote SOX2+ progenitor differentiation into metaplastic KRT5+ basal cells. Hh = hedgehog, 4OHT= 4-hydroxytamoxifen. Scale bars, 100 μm. See also Figure S3 and Table S1.

To determine whether mesenchymal Hh activation promotes the differentiation of isolated airway progenitors *in vitro*, we co-cultured Hh-activated lung mesenchyme with ITGB4+ airway progenitors from uninjured lungs in our 3D organoid assay. To activate Hh in the lung mesenchyme, we first generated a transgenic animal (*Ubc^creERT2/+^:R26^SmoM2/+^*) where constitutively active *SmoM2* could be induced in isolated lung mesenchymal cells *in vitro* utilizing a *creERT2* allele driven by a ubiquitously expressing promoter as previously described (Peng et al., 2015; Wang et al., 2018). We activated Hh in isolated lung mesenchyme by addition of 4-hydroxytamoxifen (4OHT), followed by co-culture with wild type ITGB4+ progenitors in our 3D organoid assay. The organoids are analyzed after 12-14 days in culture by separating the epithelial organoids from the mesenchyme for analysis (Figure 2C). Activation of Hh in the mesenchyme resulted in appropriate induction of *Gli1* in the separated mesenchyme (Figure 2C). But more interestingly, Hh activation in the mesenchyme significantly increased the expression of *Krt5* in the epithelial organoids and decreased the expression of *SFTPC* (Figure 2D,E). These results demonstrate that Hh activation of mesenchyme promotes the metaplastic differentiation of airway progenitors into KRT5+ cells during fibrotic repair, likely through the alteration of niche factors secreted by *Gli1*+ mesenchyme (Figure 2F).

### Single cell transcriptome analysis of *Gli1*+ mesenchyme reveals upregulated BMP antagonism in the fibrotic niche

To determine transcriptome alterations that could modify *Gli1*+ cells’ ability to alter airway progenitor behavior, we performed single cell RNA-sequencing (scRNA-seq) on sorted *Gli1* Lin+ mesenchymal cells before and after fibrotic injury (Figure 3A). 17,620 cells were captured from lineage labeled *Gli1* Lin+ cells from *Gli1^creERT/+:^R26^YFP/YFP^* lungs treated with PBS or bleomycin, with a median of 2402 genes detected per cell on the 10X Genomics platform for each condition. Library preparation was performed separately for the injured and uninjured cells, and then merged for unsupervised clustering with hierarchical analysis of gene expression. Projection of transcriptomic variations of individual cells by Uniform Manifold Approximation and Projection (UMAP) (Becht et al., 2018) shows three main clusters, defined by their anatomic localization and differentiation (Figure 3B). Based on our prior study showing that mesenchyme surrounding the proximal airway exhibits distinct identity from those in the distal alveoli (Wang et al., 2018), Cluster 1 and Cluster 2 represent the proximal and distal subsets respectively, whereas Cluster 3 represents a *de novo* population expressing myofibroblast markers that appears only during fibrotic injury in the alveoli (Figure 3C). Bleomycin-injured *Gli1* Lin+ cells demonstrate an increase in proportion of cells in the distal subset (Figure 3D), and gave rise to the entire myofibroblast subset that generates scarring in the distal alveoli along with portions of the distal subset (Figure 3D, green box). To assess potential lineage relationship between the mesenchymal subsets during fibrotic repair *in silico*, we performed RNA velocity analysis (La Manno et al., 2018) on *Gli1* Lin+ mesenchymal cells treated with bleomycin. Utilizing unbiased analysis of unspliced to spliced mRNA ratios at the single cell level to infer transcriptional trajectory, RNA velocity predicted two distinct trajectories where proximal mesenchymal cells gives rise to either distal mesenchymal cells or myofibroblasts during fibrotic repair (Figure 3E). This analysis suggests that fibrotic injury pushes proximal *Gli1* Lin+ mesenchyme towards a more distal cell fate, supported by our *in vivo* lineage tracing data showing increased abundance of *Gli1* Lin+ cells in the alveoli and differentiation into myofibroblasts localized exclusively in the alveoli after bleomycin injury (Figure S1B).

**Figure 3.**
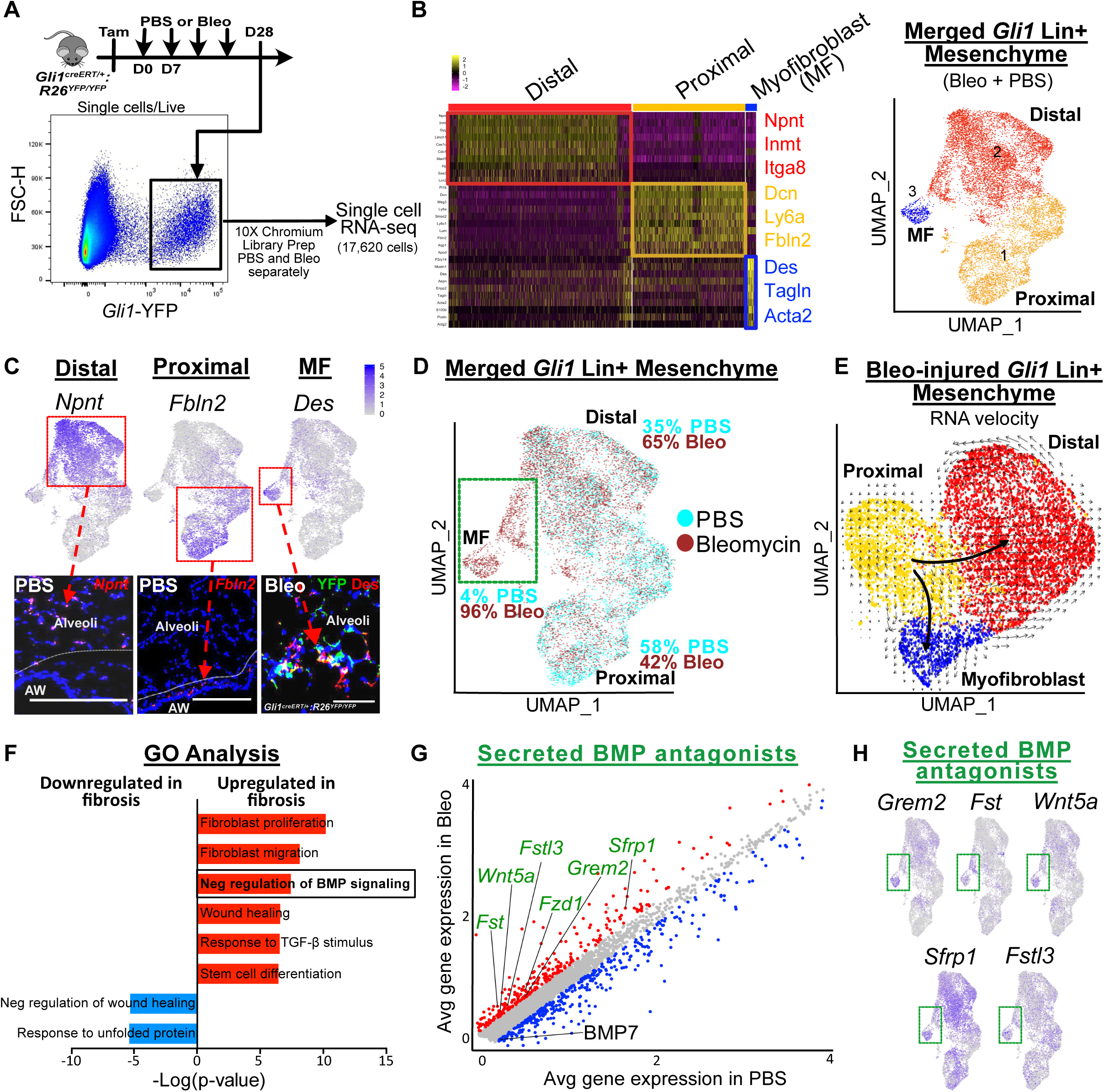
Single cell transcriptome analysis of *Gli1*+ mesenchyme reveals upregulated BMP antagonism in the fibrotic niche. (A) Workflow to isolate *Gli1* Lin+ cells during homeostasis (PBS) and fibrosis (bleomycin) for single cell analysis. (B) Left: Heatmap of merged *Gli1* Lin+ cells (PBS and bleomycin) segregates into 3 distinct clusters based on transcriptome, with highest expressing cluster-defining genes to the right. Right: UMAP plot of the three distinct clusters that appears segregated based on anatomic location. (C) *in situ* (RNAscope) and immunohistochemistry analysis shows spatial distribution of signature genes for distal, proximal, and myofibroblast clusters. (D) UMAP plot with cells of origin based on treatment shows an increase in proportion of cells in the distal and myofibroblast cluster after fibrotic injury. Portion of the distal cluster and the entire myofibroblast cluster is derived from bleo-injured cells (green box). (E) RNA velocity analysis predicting the transcriptional trajectory of *Gli1* Lin+ cellular clusters suggests proximal *Gli1*+ cell transdifferentiation into distal or myofibroblast identity during fibrotic repair. (F) GO term analysis of differentially-expressed genes between PBS and bleo in the bulk *Gli1* Lin+ cells reveals enrichment for genes involved in negative regulation of BMP signaling. (G) Gene correlation plot with each dot representing a gene, with genes significantly upregulated in fibrotic *Gli1* Lin+ cells in red (adj. p < 0.1, logfc >0.15) and downregulated (adj. p < 0.1, logfc <-0.15) in blue. Secreted BMP antagonists (GO term: negative regulation of BMP signaling) are labeled in green. (H) Gene feature plots of secreted BMP antagonists show enrichment in myofibroblast and part of distal cluster (green box). MF = myofibroblast, UMAP = Uniform Manifold Approximation and Projection, GO = Gene Ontology, BMP = Bone morphogenetic protein Scale bars, 100 μm. See also Figure S4.

To profile the transcriptional alterations in *Gli1* Lin+ mesenchyme during fibrosis, we performed differential expression analysis between *Gli1* Lin+ cells treated with PBS vs. bleomycin. Fibrotic *Gli1* Lin+ mesenchyme showed upregulation of genes classified as “TGF-β signaling components” and “negative regulation of BMP signaling” according to Gene Ontology (GO) analysis (Figure 3F,G). The upregulation of secreted BMP antagonists, particularly *Grem2* and *Fst*, is most pronounced in the *Gli1* Lin+ cells that appear *de novo* in bleomycin, including the myofibroblast cluster (Figure 3H). When we examined the expression of these secreted BMP antagonists in a publicly available cellular atlas of the murine lung (Tabula Muris et al., 2018), we see that they are almost exclusively expressed in the mesenchymal subsets of the stroma (Figure S4A). Additionally, numerous BMP ligands are also expressed in the *Gli1*+ mesenchyme, including *Bmp3, 4, 5,* and *7* (Figure S4B), although only *Bmp7* was significantly dysregulated between bleomycin and PBS (down regulated, logfc<-0.15). Our single cell analysis suggests that *Gli1* Lin+ mesenchyme upregulates local BMP antagonism during fibrotic repair through the secretion of soluble BMP antagonists that are capable of interfering with BMP ligand binding in the fibrotic niche.

### Hh activation upregulates BMP antagonism in the fibrotic lung

During organ morphogenesis, Hh and BMP signaling are often mutually antagonistic, forming antiparallel gradients of activation within the same tissue to define segmental cell fate (Zagorski et al., 2017). Activation of the BMP receptor leads to intracellular phosphorylation of SMAD1, 5, and 8, which serves as a readout for BMP activation (Massague et al., 2005). To determine the effect of Hh activation on BMP activation, we quantified the areas of the lung stained by phosphorylated-SMAD1/5/8 (pSMAD1/5/8) in the bleomycin-injured lungs. In the injured control lungs, pSMAD1/5/8 staining appears in the airway epithelium and patches of normal appearing alveoli adjacent to areas of damaged lung. Histological quantification showed a reduction of alveolar areas stained by pSMAD1/5/8 in the Hh-activated lungs compared to controls (Figure 4A,B), suggesting that Hh activation antagonizes BMP activation in fibrotic repair.

**Figure 4.**
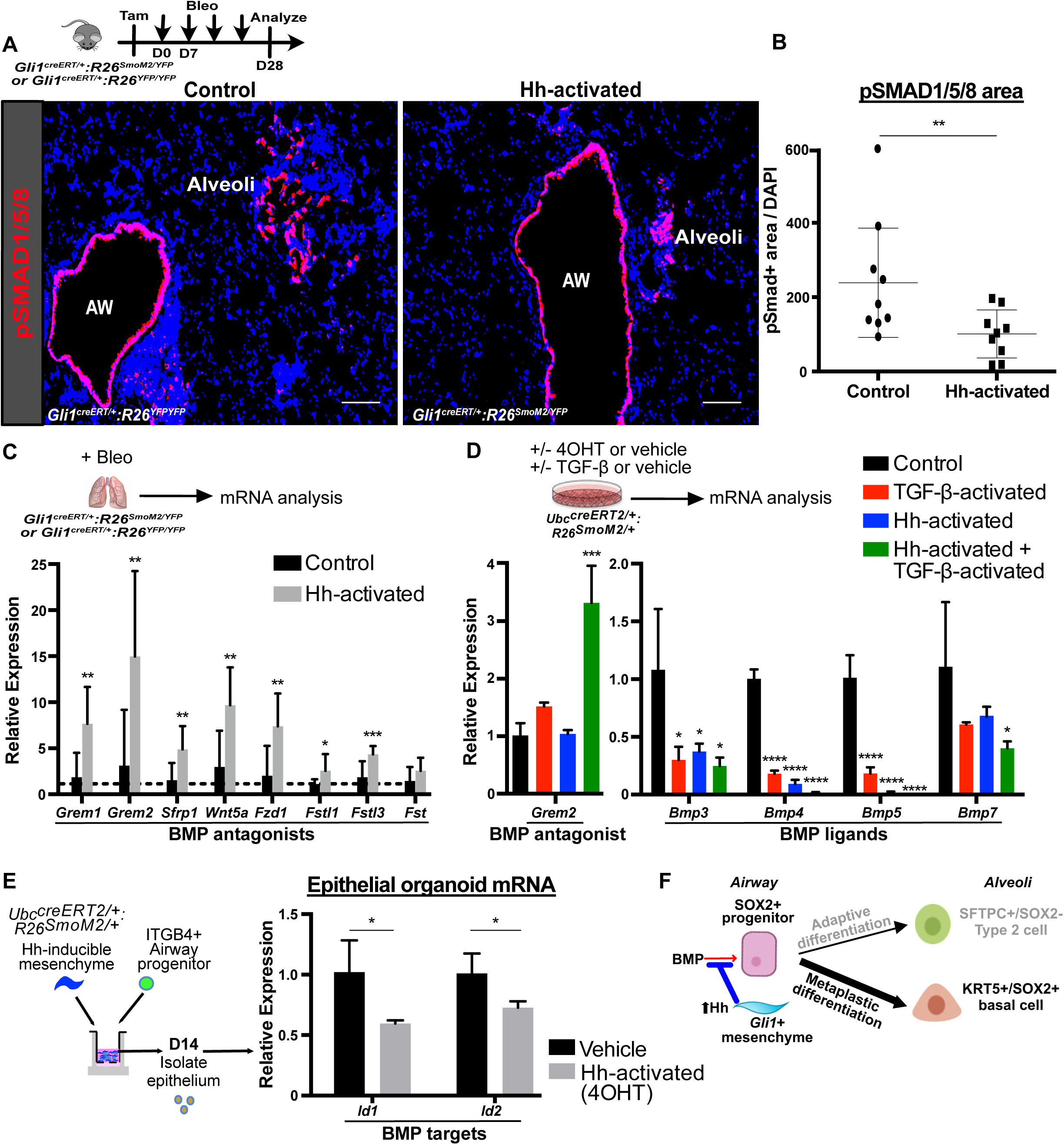
Hh activation upregulates BMP antagonism in the fibrotic lung. (A) Histology of phosphorylated-SMAD1/5/8 (pSMAD1/5/8), a marker of BMP activation, in control and Hh-activated lungs during fibrotic repair. (B) Quantification shows reduced area of pSMAD1/5/8 staining in the alveoli in the Hh-activated lungs. Data are expressed as mean ± SD. (C) qPCR analysis of whole lung mRNA shows upregulation of secreted BMP antagonists in Hh-activated lungs during fibrotic repair. Data are expressed as mean ± SD. (D) qPCR of Hh-inducible cultured mesenchyme isolated from *Ubc^creERT2/+^:R26^SmoM2/+^* lungs treated with 4OHT +/- TGF-β, shows synergistic induction of BMP antagonist *Grem2* with Hh and TGF-β activation (left), and mutually suppressive effect on BMP ligands expression (right). Data are expressed as mean ± SD. (E) qPCR of epithelial organoids isolated after co-culture with Hh-inducible mesenchyme shows reduction in BMP target genes when Hh is activated in the mesenchyme. Data are expressed as mean ± SD. (F) Model of Hh upregulating BMP antagonism in the fibrotic progenitor niche to drive metaplastic KRT5 differentiation. Scale bars, 100 μm. See also Table S1.

Next, we examined the expression of secreted BMP antagonists in the Hh-activated and control lungs treated with bleomycin. Genes encoding soluble BMP antagonists including *Grem1, Grem2, Sfrp1, Fstl1, Fstl3*, and *etc.* were all upregulated in the fibrotic Hh-activated lungs compared to controls on qPCR analysis of the whole lung (Figure 4C). These data suggest that Hh activation antagonizes BMP activation during fibrotic repair by upregulating secreted antagonists of BMP signaling in the *Gli1* Lin+ mesenchyme.

To determine whether Hh activation directly upregulates BMP antagonists *in vitro*, we collected Hh-inducible mesenchyme from the *Ubc^creERT2/+^:R26^SmoM2/+^* lungs and induced Hh activation with 4OHT. Surprisingly, Hh-activated mesenchyme showed minimal induction of the BMP antagonist, *Grem2*. Recombinant TGF-β1, a fibrotic stimulus upregulated in the activated *Gli1* Lin+ mesenchyme, also had minimal effect on *Grem2* expression when added alone to control lung mesenchyme. However, TGF-β1 activation in conjunction with Hh activation *in vitro* significantly increased the expression of *Grem2* (Figure 4D), which was the most upregulated BMP antagonists in the Hh-activated lungs *in vivo* (Figure 4C). This suggests a synergistic interaction between TGF-β and Hh in upregulating BMP antagonism in the fibrotic niche. Conversely, both Hh and TGF-β activation can independently suppress the expression of *Bmp3, Bmp4, Bmp5, and Bmp7* from the mesenchyme, with the most profound effect on *BMP4* and *BMP5* when both Hh and TGF-β are activated (Figure 4D). To determine the non-cell autonomous effect of Hh activation on airway progenitors, we isolated ITGB4+ derived epithelial organoids after co-culture with Hh-activated mesenchyme (See Figure 2C). Hh-activation in the mesenchyme reduced the expression of BMP-target genes *Id1* and *Id2* in the epithelial organoids (Figure 4E). These results demonstrate that Hh activation suppresses BMP activation by simultaneously upregulating soluble BMP antagonists in the mesenchyme and downregulating BMP ligands in the fibrotic niche, suggesting a role for BMP in directing airway progenitor fate (Figure 4F).

### BMP activation attenuates metaplastic airway progenitor differentiation

To test whether BMP activation can modify metaplastic KRT5 differentiation *in vivo*, we directly dosed recombinant BMP into the lung undergoing bleomycin treatment. Orthotopic instillation of recombinant human BMP2 (rhBMP2) is a FDA-approved therapy for spinal fusion due to its osteoinductive properties (Burkus et al., 2002). Here we chose recombinant BMP4 (rhBMP4) because *Bmp4* is expressed more highly than *Bmp2* in *Gli1* Lin+ mesenchyme (Figure S4B) and demonstrated the most profound suppression by Hh and TGF-β activation (Figure 4D). Histologically, injured wild type animals treated with rhBMP4 demonstrated a significant reduction in the areas of lung occupied by KRT5+ cystic airspace compared to vehicle (Figure 5A,B). Conversely, there is an increase in the number of SFTPC+ cells throughout the lung in the rhBMP4 treated mice (Figure 5A,B). To determine whether reversal of metaplasia would improve gas exchange in the lung, we performed oximetry on the rhBMP4 treated mice, which demonstrated significantly higher SaO2 compared to vehicle treated animals (Figure 5C). This data suggests that rhBMP4 treatment can reverse metaplasia and improve organ function in a model of lung fibrosis.

**Figure 5.**
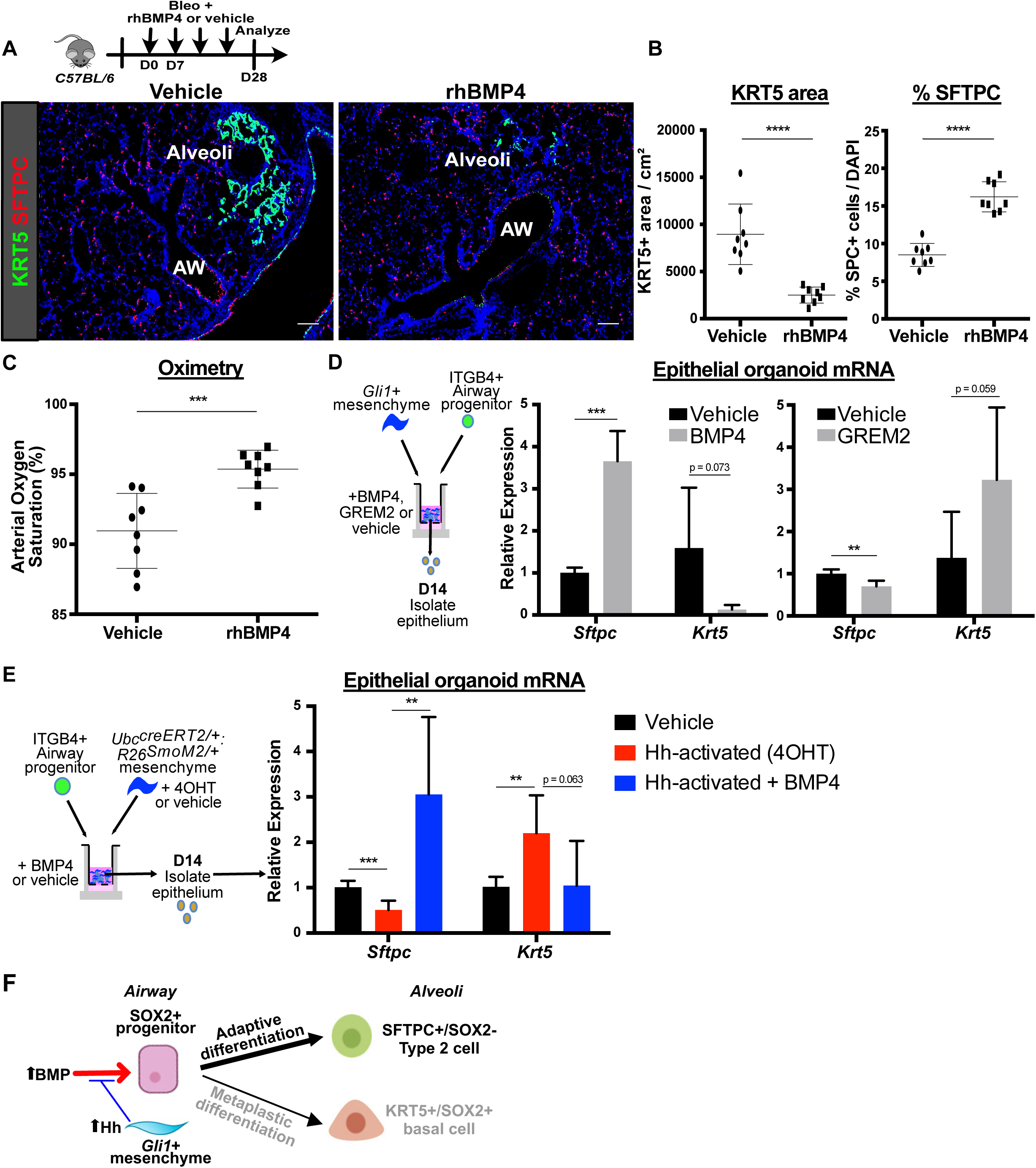
BMP activation attenuates metaplastic airway progenitor differentiation. (A) Histology shows extent of metaplastic KRT5 differentiation following treatment with intranasal inhalation of recombinant human BMP4 into wild type lungs during fibrotic injury. (B) Histologic quantification shows addition of rhBMP4 resulted in smaller areas of the lung containing KRT5+ pods and larger percentage of SFTPC+ cells in the alveoli with bleomycin injury. Data are expressed as mean ± SD. (C) Oximetry shows that animals treated with rhBMP4 had significantly improved oxygen saturation compared to vehicle-treated controls. Data are expressed as mean ± SD. (D) Left: qPCR of epithelial organoid co-cultured with *Gli1*+ mesenchyme shows that recombinant BMP4 enhances SFTPC differentiation while suppressing KRT5. Right: Conversely, the BMP antagonist GREM2 acts to suppress SFTPC differentiation while promoting KRT5. Data are expressed as mean ± SD. (E) Addition of BMP4 to organoids co-cultured with Hh-activated mesenchyme promotes SFTPC differentiation and attenuates the KRT5 differentiation induced by Hh signaling. Data are expressed as mean ± SD. (F) Model of BMP activation promoting airway progenitor differentiation into adaptive SFTPC+ Type 2 cells. AW = airway, Scale bars, 100 μm. See also Table S1.

To test the effect of BMP modulation specifically in the airway progenitor niche, we co-cultured ITGB4+ airway progenitors and *Gli1*+ mesenchyme with or without recombinant BMP4 in our 3D organoid assay. Addition of BMP4 increased the expression of *SFTPC* in the epithelial organoid and reduced the expression of *Krt5* (Figure 5D). Conversely, addition of the secreted BMP antagonist, recombinant GREM2, promoted *Krt5* expression at the expense of *SFTPC*. (Figure 5D). Furthermore, addition of BMP4 to airway progenitors co-cultured with Hh-activated mesenchyme attenuated KRT5 metaplasia in the organoids (Figure 5E), further supporting that BMP antagonism occurs downstream of Hh-activation to drive KRT5 differentiation. Overall, these data demonstrate that BMP activation promotes the differentiation of mobilized airway progenitors towards differentiation into adaptive SFTPC+ alveolar epithelium (Figure 5F), and the reversal of epithelial metaplasia is physiologically significant for improving gas exchange.

### IPF lungs display altered BMP activation in the epithelium

To determine whether fibrotic human lung mesenchyme undergoes similar transcriptomic alterations as seen in the murine *Gli1*+ mesenchyme, we analyzed single cell RNA-sequencing (scRNA-seq) of human lungs from IPF and normal controls (N=3 per group). The diseased lung was collected from a patient with an established diagnosis of IPF undergoing lung transplantation, and the control was from a sex and age-matched cadaveric donor lung without prior history of lung disease. Library preparation was again performed separately on the IPF and normal lung, and mesenchymal cells were further segregated based on the use of *Pdgfra* as a mesenchymal marker, with *Pdgfra*+ cells from each lung merged for comparison (Figure 6A). UMAP analysis shows emergence of cellular subsets in the fibrotic lung that are distinct from *Pdgfra*+ cells from healthy controls (Figure 6B). Further clustering shows a subset (Cluster 3) derived almost entirely from the IPF lung that is enriched for scar components such as genes associated with myofibroblast differentiation (*ACTA2* and *ASPN*) and matrix deposition (*COL1A1*) (Figure 6C,D). Differential gene expression with GO analysis shows a significant enrichment of BMP antagonists in the fibrotic human lung mesenchyme compared to normal controls (Figure 6E,F). Furthermore, the upregulated BMP antagonists demonstrate enrichment in the scar-forming fibroblasts (Cluster 3), especially *SFRP4* and *NBL1* (Figure 6G). This data suggests that upregulation of BMP antagonists in the scars of fibrotic lung is a conserved feature across species.

**Figure 6.**
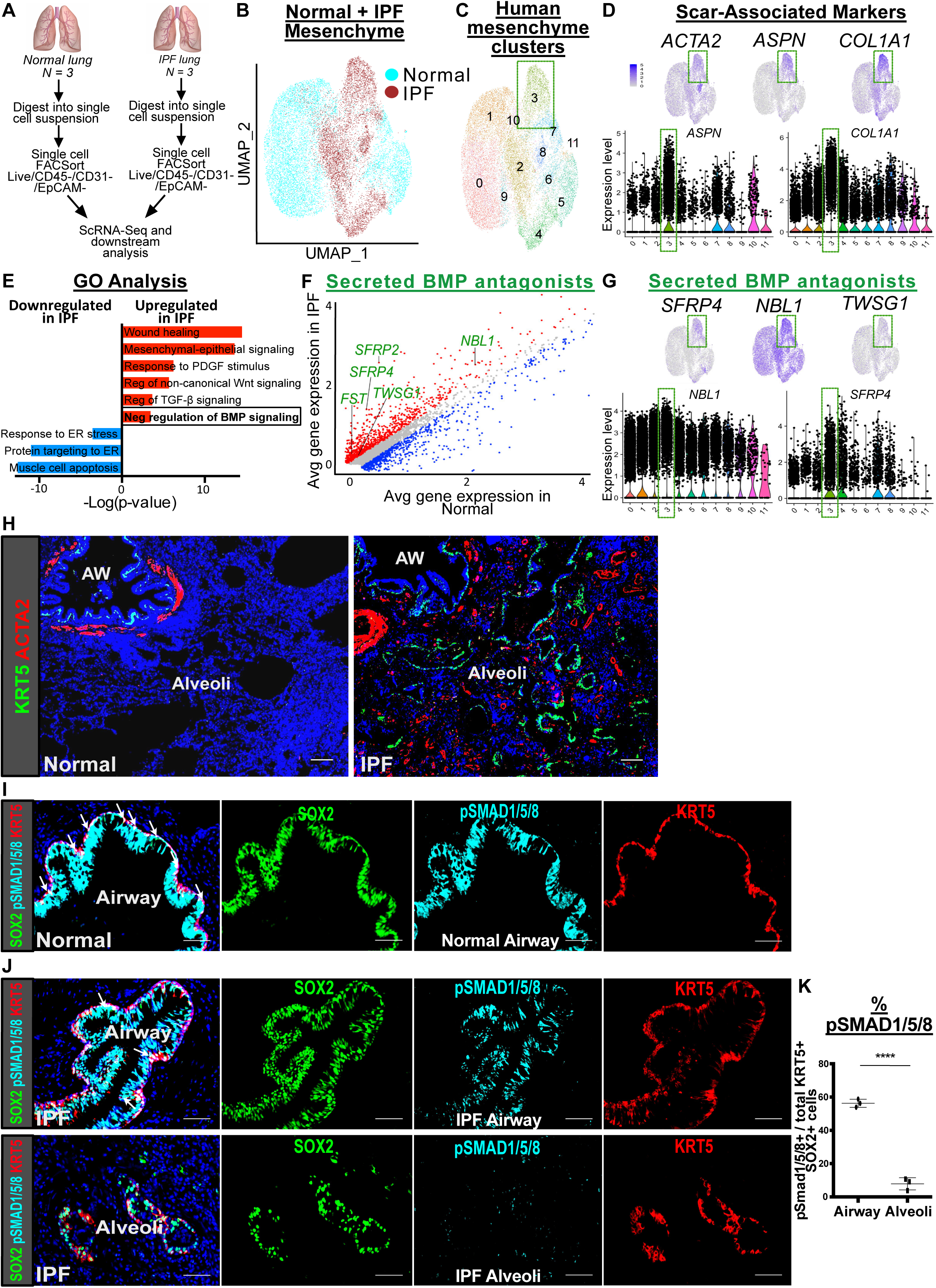
IPF lungs display altered BMP activation in the epithelium. (A) Workflow to isolate mesenchymal cells from explanted IPF lungs along with normal donor lungs for single cell analysis. (B) UMAP plot with cells of origin based on disease state (IPF vs. normal) shows distinct cellular populations that emerge in the fibrotic state. (C) UMAP plot of the distinct clusters that emerge in the normal and diseased mesenchyme, including cluster 3 that has a strong signature for scar components. (D) Gene feature plots and violin plots show that cluster 3 is highly enriched in scar-associated genes such as myofibroblast differentiation markers, *ACTA2* and *ASPN*, along with scar matrix component, *COL1A1*. (E) GO term analysis of differentially-expressed genes between IPF and normal lung mesenchyme shows enrichment for genes involved in negative regulation of BMP signaling. (F) Gene correlation plot with each dot representing a gene, with genes significantly upregulated in IPF in red (adj. p < 0.1, logfc >0.15) and downregulated (adj. p < 0.1, logfc <-0.15) in blue. Secreted BMP antagonists (GO term: negative regulation of BMP signaling) are labeled in green. (G) Gene feature plots and violin plots show that cluster 3 is also highly enriched in secreted BMP antagonists, particularly *NBL1* and *SFRP4*. (H) Histology of normal and IPF lungs shows the metaplastic presence of KRT5+ pods and ACTA2+ fibroblastic foci in the alveoli in IPF. (I) pSMAD1/5/8 staining of distal airway from normal lung shows uniform co-localization with SOX2 and KRT5 at the base of the pseudostratified epithelium. (J) Top: pSMAD1/5/8 staining of distal airway from IPF lung shows scattered co-localization with SOX2+/KRT5+ basal cells and disorganized positioning of the basal cells away from the base of the pseudostratified epithelium. Bottom: absence of pSMAD1/5/8 staining in the metaplastic SOX2+/KRT5+ basal cells in the alveoli. (K) Histologic quantification of IPF lungs shows significantly reduced pSMAD1/5/8 staining in the alveolar SOX2+/KRT5+ basal cells relative to their airway counterpart. Data are expressed as mean ± SD. Scale bars, 100 μm. See also Figure S5.

Histological analysis of IPF lungs demonstrates areas of intense scarring in the distal alveoli, marked by the presence of ACTA2+ fibroblastic foci, which are often localized adjacent to metaplastic KRT5+ cells, neither of which are present in the normal alveoli (Figure 6H). Similar to mouse, metaplastic KRT5+ basal cells in the alveoli of IPF lungs co-label with SOX2, which is normally a marker of airway differentiation in the human lung as well (Plantier et al., 2011)(Figure 6J). Cell-to-cell distance quantification of metaplastic SOX2+/KRT5+ cells in the alveoli demonstrates that they are equally proximate to the ACTA2+ fibroblastic foci when compared to the endogenous SFTPC+ epithelium (Figure S5A), suggesting a capacity for interaction between metaplastic SOX2+/KRT5+ cells and the scar.

We then examined BMP activation in the human lung using pSMAD1/5/8 staining. Unlike the mouse lung, KRT5+ basal cells extend more distally in the normal human airway, lining the basal side of the pseudostratified epithelium of the distal airways (Hogan et al., 2014). Analysis of pSMAD1/5/8 shows intense staining in the airway epithelium of normal lungs, including uniform expression in the KRT5+ basal cells (Figure 6I). However, in the IPF lungs, the airway epithelium appears dysplastic, with overgrowth of the SOX2+ epithelium and non-uniform distribution of pSMAD1/5/8 staining within the disorganized KRT5+ basal cell layer (Figure 6J). Furthermore, metaplastic SOX2+/KRT5+ basal cells in the alveoli demonstrates a significant reduction in pSMAD1/5/8 staining compared to their airway counterparts (Figure 6J,K), suggesting a down-regulation of BMP signaling in the ectopic KRT5+ cells in the alveoli, similar to what we observed in the mouse (data not shown). Taken together, these data suggest that the fibroblastic scars in IPF upregulates BMP antagonism in the local environment, concurrent with a downregulation of BMP activation in metaplastic KRT5+ cells that appears to be a conserved feature of epithelial metaplasia in lung fibrosis.

## Discussion

The ectopic appearance of mature epithelial cell types under pathological settings (*i.e.* metaplasia) is a feature of chronic wounds that often appears at tissue transition zones (Giroux and Rustgi, 2017). A very prevalent example is Barrrett’s esophagus, a common condition where chronic acid reflux leads to replacement of squamous esophageal epithelium with columnar intestinal epithelium at the gastro-esophageal junction. Despite the prevalence of metaplasia across various mucosal barrier epithelia, little is known about the processes dictating this specific form of cellular plasticity that accompanies wound repair. Another feature of wound repair is the presence of scar tissue, formed by myofibroblasts and other mesenchymal cells that enhance matrix deposition and tensile strength at the site of injury (Humphreys, 2018). While scarring and metaplasia are often found adjacent to each other, their functional relationship has not been established. Our study demonstrates how a putative MSC population (*Gli1*+ mesenchymal cells) that contributes to scarring can also promote epithelial metaplasia, providing a functional link between two commonly observed pathological phenomena in organ fibrosis. Utilizing temporally and spatially-precise *in vivo* manipulation and *ex vivo* cellular modeling, we show that *Gli1*+ mesenchyme integrates Hh activation to modify the extracellular BMP environment, which in turn regulates the metaplastic fate of adjacent airway progenitors in fibrotic repair (Figure 7).

**Figure 7.**
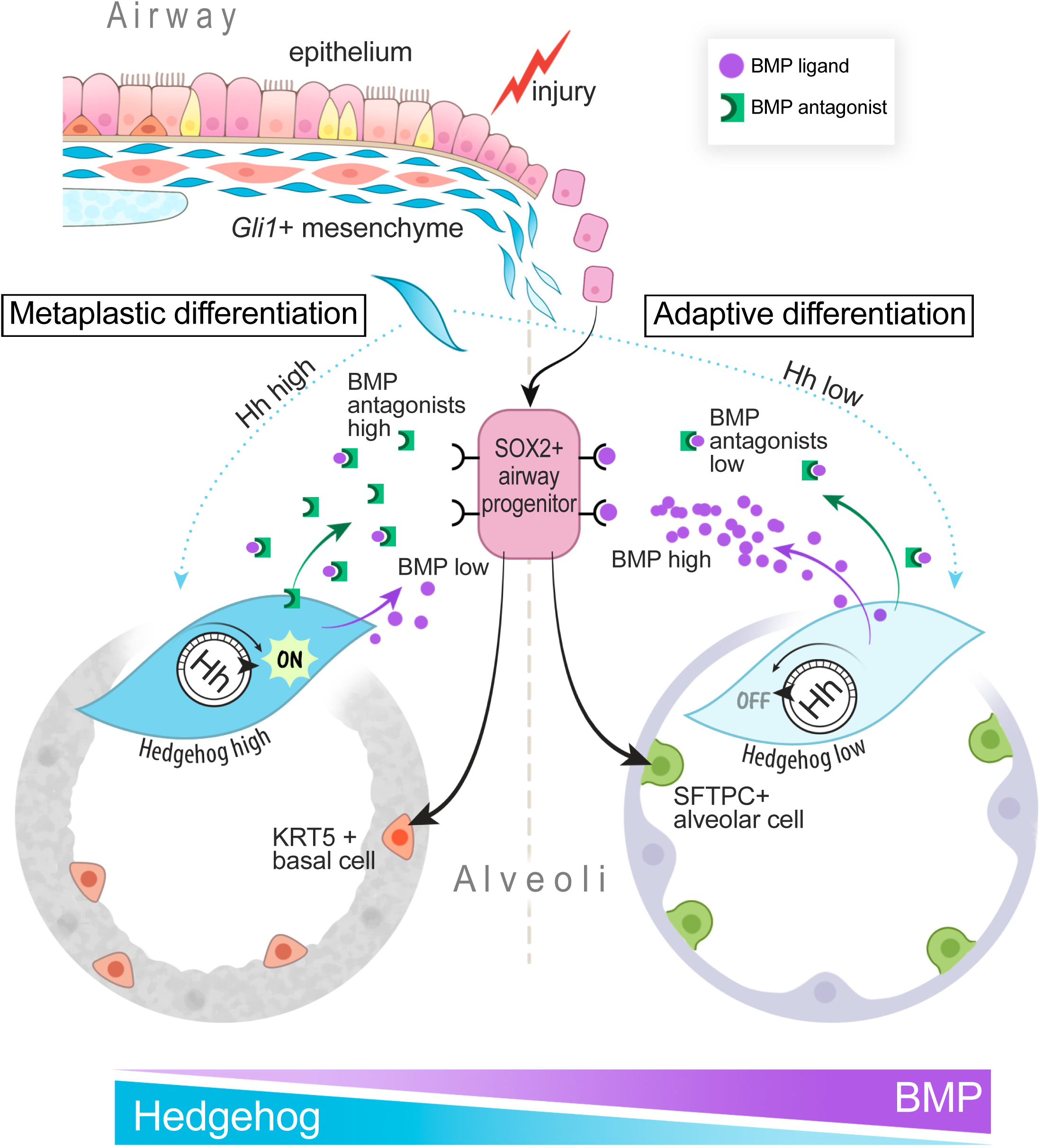
Model of *Gli1*+ mesenchyme integrating Hh activation as a rheostat that controls local BMP activation to determine SOX2+ progenitor fate.

The epithelial compartment of the proximal conducting airway contains a diverse repertoire of progenitor populations that show remarkable plasticity for interconversion *in situ* during states of homeostasis and repair (Hogan et al., 2014). Furthermore, the discovery that proximal airway progenitors can contribute to repair of the distal alveolar compartment demonstrates their incredible mobility, and raises questions about how progenitors adapt to their new local microenvironment when mobilized for tissue repair. In the setting of severe influenza infections, it has been reported that tissue hypoxia can regulate Notch/Wnt activation within airway progenitors to modify differentiation within the alveoli (Xi et al., 2017). More recently, it has been shown that FGF signaling can also alter the fate of airway progenitors after fibrotic injury (Yuan et al., 2019).

However, stem/progenitor cells do not act in isolation, and these studies raise more questions about the identity and function of the niche cells that are capable of modifying progenitor cell behavior. Anatomically, *Gli1*+ mesenchyme represents a likely interacting partner for airway progenitors due to their extreme proximity during homeostasis, as well as their ability to migrate out into the distal alveoli after fibrotic injury. Despite the problematic definition for MSCs, *Gli1*+ cells do possess trilineage differentiation potential *in vitro* as well as potential for myofibroblastic transition *in vivo (Kramann et al., 2015; Zhao et al., 2015; Zhao et al., 2014)*. Furthermore, *Gli1*+ mesenchyme has been shown to be a source of epithelial modulators in numerous organs including the lung (Degirmenci et al., 2018; Lim et al., 2014; Peng et al., 2015). Our experiments show that *Gli1*+ mesenchyme and airway progenitors form a dynamic niche during fibrotic repair, where the mesenchyme is capable of modifying local cues for progenitor fate.

Hh and BMP are key developmental morphogens that often form activation gradients in an antiparallel orientation to generate asymmetric patterning. Cells in the developing foregut and neural tube sense the local balance between Hh and BMP activation that specifies ventral (Hh predominant) vs. dorsal (BMP predominant) fate (Zagorski et al., 2017). While mutual antagonism between Hh and BMP is crucial for morphogenesis, questions remain as to whether and how the two pathways continue to interact in adult tissues during maintenance and repair. During hair follicle development and regeneration, Hh activation of the dermal papilla upregulates the expression of the secreted BMP antagonist, Noggin, which in turn suppresses BMP activation in the stem cell niche to promote follicle basal cell proliferation (Hsu et al., 2014; Woo et al., 2012). In the respiratory system, it has been reported that BMP antagonists are upregulated after injury to the tracheal epithelium, and inhibition of BMP drives basal cell proliferation (Mou et al., 2016; Tadokoro et al., 2016). Our single cell data shows that *Gli1*+ mesenchyme is the key source of numerous soluble BMP antagonists, along with BMP ligands that are expressed in the mesenchyme and other cell types. This suggests a cellular microenvironment in the adult lung where the balance between BMP ligands and soluble antagonists is tightly regulated, and Hh controls the secretion of BMP antagonists in *Gli1*+ mesenchymal cells to modulate the state of local BMP activation. We have previously shown that Hh maintains proximal mesenchymal cell identity (Wang et al., 2018), but what is surprising about our new data is that Hh also appears to promote proximal epithelial cell fate in trans by driving BMP antagonism in the fibrotic progenitor niche. This suggests a mechanism where Hh and BMP continue to compete in adult tissues to segregate compartmental identity during regeneration, by acting through a mesenchymal effector that modifies epithelial cell fate.

Over-activation of Hh has been reported in patients with IPF (Bolanos et al., 2012; Jia et al., 2017; Lee et al., 2018), but pharmacologic inhibition of the pathway has produced mixed results in experimental models of lung fibrosis (Liu et al., 2013; Moshai et al., 2014). The overwhelming endpoints of these experimental fibrotic studies have been the reduction of scarring, as measured by collagen content in the lungs as a therapeutic endpoint. However, our study demonstrates that epithelial metaplasia is an equally important endpoint in lung fibrosis, as therapeutic modifiers that target metaplasia can also reverse physiologic endpoints. We found that administration of rhBMP4 is able to promote the repletion of the alveolar epithelium with endogenous SFTPC+ type 2 cells, which also improved gas exchange as seen in the improved oximetry. The ectopic formation of basal cells in the alveoli would likely have an adverse effect on the gas exchange function of the alveoli, which would explain why their presence directly correlates with disease severity in IPF (Prasse et al., 2019). While the cellular origin of the ectopic KRT5+ basal cells in human IPF remains unclear, the downregulation of BMP activation in the ectopic alveolar KRT5+ basal cells compared to airway KRT5+ basal cells suggests that BMP could regulate the “bronchiolization” of alveoli. Our study highlights the need for other physiologically-relevant endpoints in experimental fibrosis studies, and suggest that epithelial metaplasia presents both an attractive therapeutic target as well as a clinical endpoint for tissue fibrosis.

## Author Contributions

M.C. and T.P. conceived the experiments. M.C., C.W., J.K., T.T., P.M., M.M., and P.W. performed the experiments and data analysis. D.S., A.M., and H.C. provided expertise and feedback. M.C. and T.P. wrote the manuscript.

## Acknowledgements

We thank Ka Neng Cheong, Alexis Brumwell, and Nabora Reyes de Mochel for providing technical assistance; Chris Gralapp for assistance with model illustration; Mark Looney for critical review of manuscript; the Parnassus Flow Cytometry Core for assistance with cell sorting for bulk and single cell RNA analysis (P30DK063720); Biological Imaging Development Core members (P30 DK063720); Eunice Wan and the Institute for Human Genetics Core for processing of single cell RNA samples and high-throughput sequencing. GEO accession number for raw RNA sequencing data is listed in Materials and Methods. This work is supported by NIH grants DP2AG056034, K08HL121146, R01HL142552 to T.P., along with Tobacco Related Disease Research Program New Investigator Award and Pulmonary Hypertension Association award to T.P. and Nina Ireland Program Award to M.M. for human lung collection.

## STAR★METHODS

Detailed methods are provided in the online version of this paper and include the following:

- KEY RESOURCES TABLE
- CONTACT FOR REAGENT AND RESOURCE SHARING
- EXPERIMENTAL MODEL AND SUBJECT DETAILS

- Human Lung Tissue
- Animal Studies and Treatment
- METHOD DETAILS

- Histology and Immunofluorescence
- Cleared Thick Slice Imaging
- Lung Digestion and Fluorescence Activated Cell Sorting (FACS)
- Single Cell Capture and Sequencing
- Cell Culture
- Organoid Assay
- Quantitative RT-PCR
- RNAscope *In Situ* Hybridization
- Pulse Oximetry
- QUANTIFICATION AND STATISTICAL ANALYSIS

- Statistical Analysis
- Immunofluorescence Image Quantification
- Single Cell RNA-Seq Analysis
- DATA AND SOFTWARE AVAILABILITY

## STAR★METHODS

### KEY RESOURCES TABLE

| REAGENT or RESOURCE | SOURCE | IDENTIFIER |
| --- | --- | --- |
| **Antibodies** |  |  |
| Chicken polyclonal anti-GFP (used at 1:250) | Aves Labs | Cat#GFP-1020; RRID: AB_10000240 |
| Goat polyclonal anti-GFP (used at 1:250) | Abcam | Cat#ab6673; RRID: AB_305643 |
| Rabbit polyclonal anti-ACTA2 (used at 1:200) | Abcam | Cat#ab5694; RRID: AB_2223021 |
| Rabbit polyclonal anti-Desmin (used at 1:200) | Abcam | Cat#ab15200; RRID: AB_301744 |
| Goat polyclonal anti-SCGB1A1 (used at 1:500) | Santa Cruz | Cat#sc-9772; RRID: AB_2238819 |
| Rabbit polyclonal anti-SFTPC (used at 1:2000) | Millipore Sigma | Cat#AB3786; RRID: AB_91588 |
| Goat polyclonal anti-SFTPC (used at 1:200) | Santa Cruz | Cat#sc-7706; RRID: AB_2185507 |
| Goat polyclonal anti-SFTPC (used at 1:200) | Santa Cruz | Cat#sc-7705; RRID: AB_2185505 |
| Chicken polyclonal anti-KRT5 (used at 1:500) | BioLegend | Cat#905901; RRID: AB_2565054 |
| Goat polyclonal anti-SOX2 (used at 1:200) | Neuromics | Cat#GT15098; RRID: AB_2195800 |
| Chicken polyclonal anti-RFP (used at 1:200) | Rockland | Cat#600-901-379S; RRID: AB_10703148 |
| Rabbit polyclonal anti-pSMAD1/5/8 (used at 1:100) | Millipore Sigma | Cat#AB3848-I; RRID: AB_177439 |
| AffiniPure Donkey anti-Chicken IgY (IgG) (H+L) Secondary Antibody, Alexa Fluor 488 (used at 1:250) | Jackson ImmunoResearch | Cat#703-545-155; RRID: AB_2340375 |
| Donkey anti-Rabbit IgG (H+L) Secondary Antibody, Alexa Fluor 555 (used at 1:250) | Thermo Fisher | Cat# A-31572; RRID: AB_162543 |
| Donkey anti-Goat IgG (H+L) Secondary Antibody, Alexa Fluor 647 (used at 1:250) | Thermo Fisher | Cat# A-21447; RRID: AB_2535864 |
| Rat anti-Mouse CD45 Alexa Fluor 700 (used at 1:200) | BD | Cat#560510; RRID: AB_1645208 |
| Rat anti-Mouse CD31 APC/Fire 750 (used at 1:200) | BioLegend | Cat#102528; RRID: AB_2721491 |
| Rat anti-Mouse CD326 BV421 (used at 1:200) | BD | Cat#563214; RRID: AB_2738073 |
| Rat anti-Mouse CD104 PE (used at 1:100) | BioLegend | Cat#123610; RRID: AB_2563544 |
| Biotin anti-mouse CD326 (used at 1:250) | BioLegend | Cat#118204; RRID: AB_1134178 |
| Mouse anti-human CD31 PE/Cy7 (used at 1:200) | BioLegend | Cat#303118; RRID: AB_2247932 |
| Mouse anti-human CD45 APC (used at 1:200) | BioLegend | Cat#304037; RRID: AB_2562049 |
| Mouse anti-human CD326 FITC (used at 1:200) | BioLegend | Cat#324204; RRID: AB_756078 |
| Mouse anti-human CD325a APC/Cy7 (used at 1:200) | BioLegend | Cat#349116; RRID: AB_2650978 |
| **Biological Samples** |  |  |
| Human IPF Lung Tissue | UCSF Interstitial Lung Disease Blood and Tissue Repository | N/A |
| Normal Human Lung Tissue | Northern California Transplant Donor Network | N/A |
| **Chemicals, Peptides, and Recombinant Proteins** |  |  |
| Fluorsave | Millipore Sigma | Cat#345789 |
| DAPI (used 1:1000) | Thermo Fisher | Cat#1738176 |
| DRAQ7 (used 1:1000) | Cell Signaling Technologies | Cat#7406S |
| RBC Lysis Buffer (10x) | Thermo Fisher | Cat#NC9067514 |
| Normal Donkey Serum | Thermo Fisher | Cat#50413275 |
| 4% Paraformaldehyde in PBS | Santa Cruz | Cat#sc-281692 |
| Xylene | VWR | Cat#89370-088 |
| Paraffin | Thermo Fisher | Cat#8330 |
| OCT | VWR | Cat#25608-930 |
| Diva Decloaker Antigen Retrieval (10x) | Biocare Medical | Cat#DV2004MX |
| RNAscope Target Retrieval (10x) | Advanced Cell Diagnostics | Cat#322000 |
| TritonX-100 | Sigma Aldrich | Cat#X100 |
| Tween 20 | Thermo Fisher | Cat#BP337500 |
| Histodenz | Sigma-Aldrich | Cat#D2158 |
| Sodium Azide | Sigma-Aldrich | Cat#S2002 |
| TrueBlack (50x) | Biotium | Cat#23007 |
| DMEM/F-12 | Thermo Fisher | Cat#11330032 |
| DMEM | Thermo Fisher | Cat#31053028 |
| RPMI | Millipore Sigma | Cat#R7388 |
| TrypLE Select Enzyme | Thermo Fisher | Cat#12563011 |
| Fetal Bovine Serum, heat inactivated (FBS) | Thermo Fisher | Cat#12676011 |
| Penicillin/Streptomycin (100x) | Thermo Fisher | Cat#MT30002CI |
| Bovine Serum Albumin (BSA) | Sigma Aldrich | Cat#A4503 |
| Collagenase type I | Thermo Fisher | Cat#17100017 |
| Collagenase A | Millipore Sigma | Cat#SCR136 |
| Dispase II | Thermo Fisher | Cat#17105041 |
| DNase I | Sigma Aldrich | Cat#DN25 |
| Matrigel | Thermo Fisher | Cat#CB-40230A |
| Tamoxifen | Sigma-Aldrich | Cat#T5648 |
| 4-OH-tamoxifen (4OHT) | Millipore Sigma | Cat#579002 |
| Corn oil | Sigma Aldrich | Cat#C8267 |
| FcR Blocking Reagent | Miltenyi Biotec | Cat#130-092-575 |
| OneComp Beads | Thermo Fisher | Cat#501129031 |
| SuperScript III Reverse Transcriptase | Thermo Fisher | Cat#18080044 |
| RnaseOUT Rnase Inhibitor | Thermo Fisher | Cat#100000840 |
| Thiazovivin (ROCK inhibitor) | STEMCELL Technologies | Cat#72252 |
| Cholera toxin | Sigma Aldrich | Cat#C8052 |
| Trizol | Thermo Fisher | Cat#15596018 |
| Bleomycin | Hospira | N/A |
| Recombinant mouse GREM2 Protein | R&D Systems | Cat#2069-PR-050 |
| Recombinant human BMP4 Protein | R&D Systems | Cat#314-BP-010 |
| Recombinant human TGF-β1 | PeproTech | Cat#100-21 |
| Critical Commercial Assays |  |  |
| PicoPure RNA Isolation Kit | Applied Biosystems | Cat#KIT0204 |
| RNeasy Micro Kit | QIAGEN | Cat#74004 |
| E.Z.N.A Total RNA Kit | Omega Bio-Tek | Cat#R6834-01 |
| DyNAmo Flash SYBR Green qPCR Kit | Thermo Fisher | Cat#F415L |
| SAGM Bullet Kit | Lonza | Cat#CC-3118 |
| RNase-Free DNase Set | QIAGEN | Cat#79254 |
| GoTaq genotyping assay | Promega | Cat#M7823 |
| RNAscope Multiplex Fluorescent Reagent Kit v2 | Advanced Cell Diagnostics | Cat#323100 |
| RNAscope Probe Mm-Npnt | Advanced Cell Diagnostics | Cat#316771 |
| RNAscope Probe Mm-Fbln2 | Advanced Cell Diagnostics | Cat#447931 |
| TSA Plus Cyanine 3 | Perkin Elmer | Cat #NEL744001KT |
| EasySep Mouse Streptavidin Isolation Kit | STEMCELL Technologies | Cat#19860 |
| MouseOx Pulse Oximeter | STARR Life Sciences | https://www.starrrlifesciences.com/product-category/pulse-oximetry/ |
| Deposited Data |  |  |
| Murine *GLI1*+ cell scRNA-seq (PBS and Bleo) | This paper | GEO: GSE140032 |
| IPF and Normal Human Lung scRNA-seq | This paper | GEO: GSE132771 |
| Mus musculus Lung scRNA-seq | Tabula Muris | https://tabula-muris.ds.czbiohub.org/ |
| Experimental Models: Organisms/Strains |  |  |
| Mouse: C57BL/6 | Charles River Laboratories | Strain No: 027; RRID: IMSR_CRL:027 |
| Mouse: *Gli1*<sup>EGFP</sup> | GENSAT | Stock No: DM197; RRID: MMRRC_010441-UCD |
| Mouse: *Gli1*<sup>creERT</sup> | The Jackson Laboratory | Stock No: 007913; RRID: IMSR_JAX:007913 |
| Mouse: *Ubc*<sup>creERT2</sup> | The Jackson Laboratory | Stock No: 007001; RRID: IMSR_JAX:007001 |
| Mouse: *Sox2*<sup>creERT2</sup> | The Jackson Laboratory | Stock No: 017593; RRID: IMSR_JAX:017593 |
| Mouse: *R26*<sup>SmoM2</sup> | The Jackson Laboratory | Stock No: 005130; RRID: IMSR_JAX:005130 |
| Mouse: *R26*<sup>EYFP</sup> | The Jackson Laboratory | Stock No: 006148; RRID: IMSR_JAX:006148 |
| Mouse: *R26*<sup>tdTomato</sup> | The Jackson Laboratory | Stock No: 007909; RRID: IMSR_JAX:007909 |
| Mouse: *R26*<sup>mTmG</sup> | The Jackson Laboratory | Stock No: 007576; RRID: IMSR_JAX:007576 |
| Mouse: *R26*<sup>DTA</sup> | The Jackson Laboratory | Stock No: 009669; RRID: IMSR_JAX:009669 |
| Oligonucleotides |  |  |
| Primers for qPCR: listed in Table S1 | This paper; Primerbank; (Vaughan et al., 2015; Wang et al., 2018) | See Table S1 |
| **Software and Algorithms** |  |  |
| Fiji | NIH | https://www.imagej.net/Fiji |
| FlowJo | FLOWJO LLC | https://www.flowjo.com/ |
| Prism 6.0 | GraphPad Software | https://www.graphpad.com/scientific-software/prism/ |
| CellRanger v2.1.1 | 10X Genomics | https://www.support.10xgenomics.com/single-cell-gene-expression/software/pipelines/latest/installation |
| Seurat R Package | (Butler et al., 2018) | https://satijalab.org/seurat/ |
| RNA Velocity | (La Manno et al., 2018) | http://velocityto.org/ |
| GO Enrichment Analysis | (Ashburner et al., 2000) | http://geneontology.org/ |
| Imaris 8.3.1 | BitPlane | http://imaris.oxinst.com/ |

### CONTACT FOR REAGENT AND RESOURCE SHARING

Further information and requests for reagents may be directed to, and will be fulfilled by, the Lead Contact, Tien Peng.

### EXPERIMENTAL MODEL AND SUBJECT DETAILS

#### Human Lung Tissue

Studies involving human tissue were approved by the UCSF Institutional Review Board. IPF lungs were obtained from patients diagnosed with interstitial pneumonia or scleroderma at the time of lung transplant. All subjects provided written informed consent. Normal lungs were obtained from brain-dead donors that were rejected for lung transplantation.

#### Animal Studies and Treatment

All animals were housed and treated in accordance with the Institutional Animal Care and Use Committee (IACUC) protocol approved at the University of California, San Francisco. Generation and genotyping of the *Gli1^EGFP^*, *Gli1^creERT^*, *Ubc^creERT2^*, *Sox2^creERT^*, *R26^SmoM2^*, *R26^EYFP^*, *R26^tdTomato^*, *R26^mTmG^*, and *R26^DTA^* lines were performed as previously described by The Jackson Laboratory. 8-12 week old littermate mice were gender balanced and randomly assigned to experimental groups. A minimum of three mice were used per group.

For lineage tracing studies, tamoxifen (Cat#T5648; Sigma-Aldrich) was dissolved in corn oil and administered intraperitoneally at 200 mg/kg body weight per day for five consecutive days, follow by at least 7 days before bleomycin was administered. For bleomycin injury, mice were given pharmaceutical grade bleomycin (Hospira) dissolved in PBS via intranasal installation once a week for four weeks. Mixed background mice were given a dose of 1 U/kg and C57BL/6 background mice were given 0.75 U/kg per dose. Mice were sacrificed seven days after the final treatment. For BMP4 treatment, mice were treated with 5 µg/mL of human recombinant BMP4 protein (Cat#314-BP-010; R&D Systems) via intranasal installation twice a week for four weeks, concurrently with bleomycin injury. Mice were sacrificed seven days after the final bleomycin treatment and two days after the final BMP4 treatment.

### METHOD DETAILS

#### Histology and Immunofluorescence

The right ventricle of the mice were perfused with 1-3 mL PBS and the lungs were inflated with 1-3 mL 4% paraformaldehyde (PFA) in PBS, and then fixed in 4% PFA overnight at 4 °C. After fixation, the lungs were washed with cold PBS 4 times for 30 minutes each at 4 °C, and then dehydrated in a series of increasing ethanol concentration washes (30%, 50%, 70%, 95% and 100%) for a minimum of 2 hours per wash. The dehydrated lungs were incubated with xylene for 1 hr at RT and then with paraffin at 65 °C for 90 minutes twice, and then embedded in paraffin. The lungs were sectioned at 8 µm on a microtome.

For histologic analysis of organoid assays, transwells were fixed in 4% PFA overnight at 4 °C, then washed in PBS overnight at least three times. Transwells were cut and embedded in OCT, then 8 µm sections were cut on a cryostat.

For immunofluorescent staining, paraffin sections were incubated in xylene for 10 minutes twice, then rehydrated in ethanol washes (100% twice, 95%, 70%, 50% ethanol) for 5 minutes each. OCT embedded slides were fixed in 4% PFA at RT for 10 minutes, then washed three times with PBS. For both paraffin and OCT embedded slides, antigen retrieval (Cat#DV2004MX; Biocare Medical; Cat#322000; Advanced Cell Diagnostics) was performed for 30 minutes at 95 °C. Slides were washed with 0.1% Tween 20 in PBS (PBST) 3 times for 5 minutes. Slides were then incubated in blocking buffer (3% donkey serum in PBST) for at least 1 hour, then incubated overnight in primary antibodies in 1% donkey serum in PBST at 4 °C. The following primary antibodies were used for mouse tissue: chicken anti-GFP (Cat#GFP-1020; Aves Labs; used 1:1250), goat anti-GFP (Cat#ab6673, Abcam; used 1:250), rabbit anti-ACTA2 (Cat#ab5694; Abcam; used 1:200), rabbit anti-DES (Cat#ab15200; Abcam; used 1:200), goat anti-SCGB1a1 (Cat#sc-9772; Santa Cruz; used 1:500), rabbit anti-SFTPC (Cat#AB3786; Millipore Sigma; used 1:2000), goat anti-SFTPC (Cat#sc-7706; Santa Cruz; used at 1:200), chicken anti-KRT5 (Cat#905901; BioLegend; used at 1:500), goat anti-SOX2 (Cat#GT15098; Neuromics; used 1:200), chicken anti-RFP (Cat#600-901-379S; Rockland; used 1:200), and anti-p-SMAD1/5/8 (Cat#AB3848-I; Millipore Sigma; used 1:100). The following primary antibodies were used for human tissue: rabbit anti-ACTA2 (Cat#ab5694; Abcam; used 1:200), rabbit anti-p-SMAD1/5/8 (Cat#AB3848-I; Millipore Sigma; used 1:100), chicken anti-KRT5 (Cat#905901; BioLegend; used at 1:500), goat anti-SOX2 (Cat#GT15098; Neuromics; used 1:200), and goat anti-SFTPC (Cat#sc-7705; Santa Cruz; used at 1:200). Slides were washed with PBST and then incubated for 1 hour at RT in secondary antibodies diluted in PBST. The following secondary antibodies were used at 1:250: Donkey anti-Chicken IgG (H+L) Alexa Flour 488 (Cat#703-545-155; Jackson ImmunoResearch), Donkey anti-Rabbit IgG Alexa Flour 555 (Cat# A-31572; Thermo Fisher), and Donkey anti-Goat Alexa Flour 647 (Cat# A-21447; Thermo Fisher). DAPI (0.2 µg/mL) (Cat#1738176; Thermo Fisher) was added for 5 minutes, and slides were then mounted with TrueBlack to reduce background and Fluorosave to maintain fluorescence.

#### Cleared Thick Slice Imaging

Mouse lung was extracted as described above and fixed in 4% PFA overnight at 4 °C. Tissue was washed with changes of PBS for 2 hours and cut into 200 µm sections on a vibratome. Sections were blocked for at least 1 hour in 0.3% Triton X, 5% FBS, 0.5% BSA in PBS with sodium azide at 4 °C. Sections were then incubated in primary antibodies (rabbit anti-SFTPC; Cat#AB3786; Millipore Sigma; chicken anti-KRT5; Cat#905901; BioLegend; goat anti-SOX2; Cat#GT15098; Neuromics) in PBS with 0.15% Triton X, 7.5% FBS, and 0.75% BSA for 1-3 days at 4 °C. Sections were washed with 0.15% Triton X and incubated in secondary antibodies (Donkey anti-Chicken IgG (H+L) Alexa Flour 488; Cat#703-545-155; Jackson ImmunoResearch; Donkey anti-Rabbit IgG Alexa Flour 555; Cat# A-31572; Thermo Fisher; Donkey anti-Goat Alexa Flour 647; Cat# A-21447; Thermo Fisher) overnight at 4 °C. After several washes with 0.15% Triton X, DAPI was added in PBS for 30 minutes and sections were rinsed again. Sections were cleared using RIMS (40 g Histodenz, 30 mL PBS, 5 µL Tween 20, 50 µL 10% sodium azide) for 30-60 min and mounted. Images were taken on a Nikon A1R multi-photon confocal microscope.

#### Lung digestion and Fluorescence Activated Cell Sorting (FACS)

For mouse, whole lung was dissected from adult animals and tracheally perfused with a digestion cocktail of Collagenase Type I (Cat#17100017; Thermo Fisher; used 225 U/mL), Dispase II (Cat#17105041; Thermo fisher; used 15 U/mL) and Dnase I (Cat#DN25; Sigma-Aldrich; used 50 U/mL) and removed from the chest. The lung was further diced with a razor blade and incubated in digestion cocktail for 45 mins at 37 °C with continuous shaking. The mixture was then washed with sorting buffer (2% FBS and 1% Penicillin-Streptomycin in DMEM (Cat#31053028; Thermo Fisher)). The mixture was passed through a 70 µm cell strainer and resuspended in RBC lysis buffer (Cat#NC9067514; Thermo Fisher), then passed through a 40 µm cell strainer. Cell suspensions were incubated with the appropriate conjugated antibodies in sorting buffer for 30 min at 4 °C and washed with sorting buffer. Doublets and dead cells were excluded based on forward and side scatter and DRAQ7 (Cat#7406S; Cell Signaling Technologies) fluorescence, respectively. Immune and endothelial cells were excluded using CD45 (Alexa Flour 700; Cat#560510; BD; used 1:200) and CD31 (APC/Fire750; Cat#102528; BioLegend; used 1:200), respectively. For *Gli1^EGFP^* sorting, epithelial cells were also excluded using CD326 (BV421; Cat#563214; BD; used 1:200), then *Gli1* cells were sorted using endogenous GFP fluorescence. For ITGB4 sorting, cells were selected for CD326 and then CD104 (PE; Cat#123610; BioLegend; used 1:100). Cells were sorted into sorting buffer. Analysis was performed using FlowJo software.

For human, multiple pieces were collected from whole lungs and separated into smaller pieces for a total of approximately 1 gram. Tissue was cut with scissors and resuspended in a digestion cocktail of 0.25% Collagenase A (Cat#SCR136; Millipore Sigma), Dispase II (Cat#17105041; Thermo fisher; used 15 U/mL) and Dnase I (Cat#DN25; Sigma-Aldrich; used 50 U/mL) in RPMI (Cat#R7388; Millipore Sigma). The tissue was digested at 37 °C for 1 hour with intermittent resuspension. Cells were passed through a 100 µm cell strainer, washed with PBS, and resuspended in 0.5% BSA in PBS. Single cells were selected and mesenchymal cells were sorted using the following gating strategy: DAPI-, CD31-(PE/Cy7; Cat#303118; BioLegend; used 1:200), CD45- (APC; Cat#304037; BioLegend; used 1:200), CD326- (FITC; Cat#324204; BioLegend; used 1:200), and CD235a- (APC/Cy7; Cat#349116; BioLegend; used 1:200).

#### Single Cell Capture and Sequencing

For mouse scRNA-seq, all live *Gli1* Lin+ cells were sorted from two *Gli1^creERT/+^:R26^YFP/YFP^* adult lungs. *Gli1* Lin+ cells were isolated based on forward and side scatter, DAPI exclusion, and GFP fluorescence. One mouse was challenged with repetitive bleomycin and one was treated with PBS at the same timepoints as a control. The sorted *Gli1* Lin+ cells from each animal were separately resuspended in 50 µL PBS with 0.04% BSA at 1,000 cells/µL. For human scRNA-seq, sorted (live, CD31-, CD45-, CD326-, CD235a-) healthy and IPF mesenchymal cells were resuspended in 0.05% BSA in PBS. Cells were loaded onto a single lane per sample into the Chromium^TM^ Controller to produce gel bead-in emulsions (GEMs). GEMs underwent reverse transcription for RNA barcoding and cDNA amplification, with the library prepped using the Chromium Single Cell 3’ Reagent Version 2 kit. Each sample was sequenced in 1 lane of the HiSeq2500 (Illumina) in Rapid Run Mode.

#### Cell Culture

For *Ubc^creERT2/+^:R26^SmoM2/+^* mesenchymal cells, lung was digested as above and then cultured on gelatin-treated tissue culture plates in DMEM/F-12 (Cat#11330032; Thermo Fisher) with 10% FBS and 1% Penicillin-Streptomycin. Media was refreshed every other day and primary lung mesenchymal cells were maintained for no more than five passages. The *Ubc^creERT2/+^:R26^SmoM2/+^* mesenchymal cells were treated with vehicle or 1 µg/ml 4-OH-tamoxifen (4OHT) (Cat#579002; Millipore Sigma) for 72 hrs. When used for organoid assays, 4OHT treatment occurred for 72 hours immediately prior to co-culture with epithelium. Where applicable, cells were treated with vehicle or 4 ng/mL TGF-β1 (Cat#100-21; PeproTech) for 72 hours.

#### Organoid Assay

*Gli1^EGFP^* adult mouse lungs were FACS sorted for ITGB4+ epithelial progenitor cells and *Gli1*+ mesenchymal cells. ITGB4+ epithelial cells and *Gli1* mesenchymal cells were co-cultured (15×10^3^ epithelial cells : 5×10^4^ mesenchymal cells/well) in a modified MTEC media diluted 1:1 in growth factor reduced matrigel (Cat#CB-40230A; Thermo Fisher). Modified MTEC culture media (Cat#CC-3118; Lonza) is comprised of small airway basal media (SABM) with selected components from SAGM bullet kit including Insulin, Transferrin, Bovine Pituitary Extract, Retinoic Acid, and human Epidermal Growth Factor. 0.1 µg/mL cholera toxin (Cat#C8052; Sigma Aldrich), 5% FBS, and 1% Penicillin-Streptomycin were also added to the media. Cell suspension-matrigel mixture was placed in a transwell and incubated in growth media with 10 µM ROCK inhibitor (Cat#72252; STEMCELL Technologies) in a 24 well plate for 48 hours, after which the media was replenished every other day (lacking ROCK inhibitor). Each experimental condition was performed in triplicates. Where applicable, recombinant BMP4 (Cat#314-BP-010; R&D Systems; used 50 ng/mL) and GREM2 (Cat#2069-PR-050; R&D Systems; used 1.5 µg/mL) were added to the media after 48 hours and replenished in every media change. Colonies were assayed after 12-14 days.

To extract RNA from the organoid assays, cell suspension-matrigel mixtures in the transwells were washed with PBS and incubated in a digestion cocktail of Collagenase Type I (Cat#17100017; Thermo Fisher; used 225 U/mL), Dispase II (Cat#17105041; Thermo fisher; used 15 U/mL) and Dnase I (Cat#DN25; Sigma-Aldrich; used 50 U/mL) for 1 hour at 37 °C with intermittent resuspension. The mixture was removed from the transwell and resuspended in TrypLE (Cat#12563011; Thermo Fisher) and shaken at 37 °C for 20 minutes. Cells were resuspended in sorting buffer (2% FBS and 1% Penicillin-Streptomycin in DMEM) and blocked with rat serum (Cat#19860; STEMCELL Technologies; used 1:50) for 10 minutes at 4 °C. Cells were then stained with biotin anti-mouse CD326 (Cat#118204; BioLegend; 1:250) for 30 minutes at 4 °C. Streptavidin beads (Cat#19860; STEMCELL Technologies; used 1:50) were added to isolate the epithelial cells.

#### Quantitative RT-PCR

Total RNA was obtained from cells isolated from organoid assays or cultured primary lung fibroblasts using the PicoPure RNA Isolation Kit (Cat#KIT0204; Applied Biosystems) or the RNeasy Kit (Cat#74004; QIAGEN), following the manufacturers’ protocols. RNA from mouse lung tissue was obtained by removing the entire left lobe, homogenizing in trizol (Cat#15596018; Thermo Fisher), and extracting RNA using the E.Z.N.A Total RNA Kit (Cat#R6834-01; Omega Bio-Tek) following manufacturer instructions. cDNA was synthesized from total RNA using the SuperScript Strand Synthesis System (Cat#18080044, Cat#100000840; Thermo Fisher). Quantitative PCR was performed using the SYBR Green system (Cat#F415L; Thermo Fisher). Primers are listed in Table S1. Relative gene expression levels after qRT-PCR were defined using the ΔΔCt method and normalizing to GAPDH. To calculate the ratios of fold changes between KRT5 and SFTPC, the relative expression of each gene was separately calculated and normalized to GAPDH. The fold changes of each gene per sample were then divided by eachother. There were a minimum of three biological replicates for each genotype/condition.

#### RNAscope *In Situ* Hybridization

Paraffin-embedded lung sections were processed for RNA *in situ* detection of *Fbln2* (Cat#447931; Advanced Cell Diagnostics) and *Npnt* (Cat#316771; Advanced Cell Diagnostics) using the RNAscope Multiplex Fluorescent Reagent Kit v2 (Cat#323100; Advanced Cell Diagnostics) according to the manufacturer’s instructions.

#### Pulse Oximetry

The MouseOx Pulse Oximeter system (STARR Life Sciences) was used to measure arterial oxygen saturation from awake mice on the day of sacrifice. Mice were shaven prior to measurement. Mice were measured at 5 measurements per second for at least 5 minutes and at least 10 successful readings. All successful measurements (error code = 0) were averaged for each mouse.

### QUANTIFICATION AND STATISTICAL ANALYSIS

#### Statistical Analysis

All statistical analyses were performed in GraphPad Prism 6.0. Unpaired one-tailed t-tests were used to determine the p-value and data in graphs are presented as mean ± SD.

#### Immunofluorescence Image Quantification

Sections were imaged for quantification on a Zeiss Lumar V12 microscope. At least three samples per genotype/condition were used, and at least 4 randomly selected sections were chosen for each sample. Cell counts for SFTPC cells were performed on Fiji using the “Cell Counter” plug-in. Cell counts for DAPI were performed automatically by Fiji software using the “Analyze Particles” tool. For quantification by area, the image was first converted to 8 bit and the “Measure” function was used with a consistent threshold set to a minimum of 20 υm^2^ and maximum of infinity. Distance quantification was performed using Imaris 8.3.1 software. At least three samples per condition were used, and at least 6 randomly selected sections were chosen for each sample. Surfaces were created for each marker of interest, and distance between two surfaces was calculated using the “Big Sortomato” tool. The distances between all surfaces in one image were averaged. For all analyses, performer was blinded to the specimen genotype and condition. Results were averaged between each specimen and standard deviations were calculated per genotype/condition.

#### Single Cell RNA-Seq Analysis

To build transcript profiles of individual cells the CellRanger version 2.1.1 software with default settings was used for de-multiplexing, aligning reads with STAR software to GRCm38 for mouse genome and Hg19 for human, and counting unique molecular identifiers (UMIs). We used the Seurat R package along with a gene-barcode matrix provided by CellRanger for downstream analysis. In total, we filtered the data in 2 different steps. We first filtered the dataset by only accepting cells that expressed a minimum of 200 genes and genes that were expressed in at least 3 cells. Our second filter was set to accept cells with less than 6000 unique gene counts and 5% mitochondrial counts. After removing unwanted cells, we used “LogNormalize” to normalize the gene expression measurements for each cell. We calculated 2,000 features that exhibit high cell-to-cell variation, which were used in principle component analysis (PCA) after scaling the data. We used the JackStrawPlot function in the Seurat package to create Scree plots and compare p-value (significance) for each principle component. We selected 10 different PCA’s for clustering of both mouse and human cells. Clustering results were visualized using the Uniform Manifold Approximation and Projection (UMAP) algorithm in the Seurat package. For mouse datasets, we further integrated the PDGFRA+ clusters from both PBS and bleo samples to compare the datasets. Differentially expressed genes between PBS and bleo datasets were identified using a MAST test. RNA velocity was calculated on the integrated object using Velocyto package and RNA velocity in SeuratWrappers package in R. For human scRNA-seq analysis, we similarly integrated the PDGFRA+ clusters from both 3 healthy controls and 3 patients with IPF, and the differentially expressed genes were identified with a MAST test. For cluster visualization and individual gene visualization on all clusters we used the UMAP function. Gene ontology enrichment analysis was performed using the PANTHER Overrepresentation test and entering the top differentially expressed genes with adj. p <0.1 and logfc >0.15 or <-0.15.

### DATA AND SOFTWARE AVAILABILITY

The mouse sequencing data reported in this paper is deposited in NCBI Gene Expression Omnibus (GEO) under the accession number: GSE140032. The human sequencing data reported in this paper is deposited in NCBI Gene Expression Omnibus (GEO) under the accession number: GSE132771.

