## Supplemental Figures and Table for "*Gli1*+ mesenchymal stromal cells modulate epithelial metaplasia in lung fibrosis"

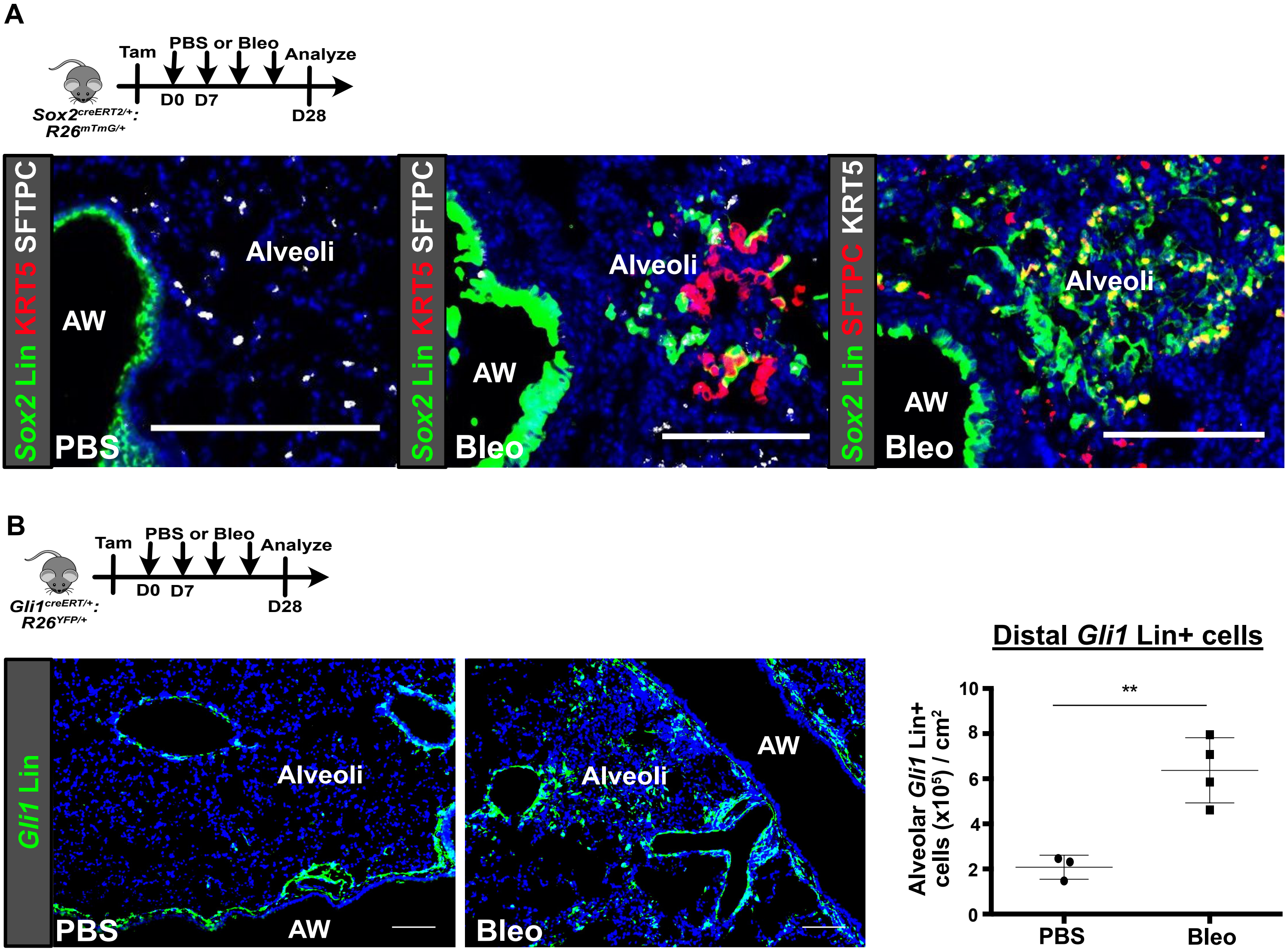
A
Tam
PBS or Bleo
Analyze
D0
D7
D28
Sox2creERT2/+:
R26mTmG/+
Sox2 Lin KRT5 SFTPC
AW
Alveoli
PBS
Sox2 Lin KRT5 SFTPC
AW
Alveoli
Bleo
Sox2 Lin SFTPC KRT5
AW
Alveoli
Bleo
B
Tam
PBS or Bleo
Analyze
D0
D7
D28
Gli1creERT/+:
R26YFP/+
Gli1 Lin
Alveoli
PBS
AW
Alveoli
AW
Bleo
Distal Gli1 Lin+ cells
Alveolar Gli1 Lin+ cells (x10^5) / cm^2
**
0
2
4
6
8
10
PBS
Bleo

**Figure S1. Epithelial progenitors and *Gli1*+ mesenchyme expand distally in fibrotic repair, related to Figure 1.**

(A) *Sox2* Lin+ cells are restricted to the airway epithelium in homeostasis. After fibrotic injury, these cells migrate into the alveoli and differentiate into either KRT5+ basal cells or SFTPC+ alveolar cells.

(B) *Gli1* Lin+ cells are located within the basement membrane of the airway during homeostasis but expand into the distal alveoli during fibrotic repair.

AW = airway. Scale bars, 100 μm.

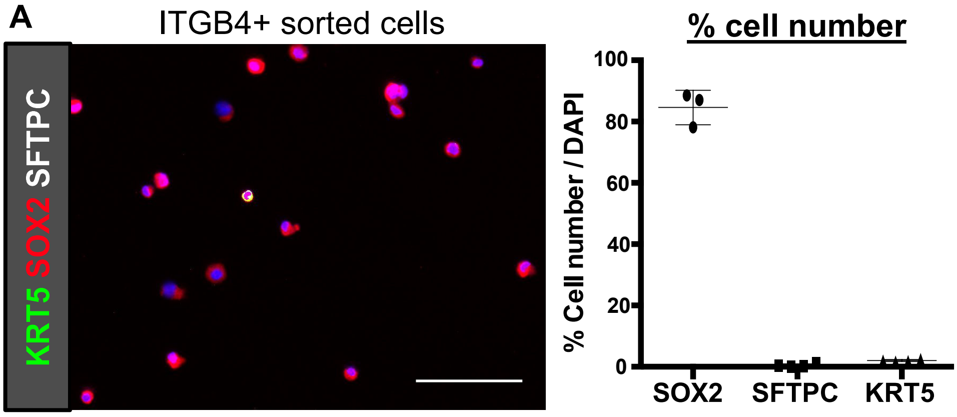
A
ITGB4+ sorted cells
KRT5 SOX2 SFTPC
% cell number
% Cell number / DAPI
100
80
60
40
20
0
SOX2
SFTPC
KRT5

**Figure S2. Sorted ITGB4+ epithelial cells are mostly SOX2+, related to Figure 1.**

(A) Cytospin of freshly sorted ITGB4+ confirms that majority of cells are SOX2+, and cells rarely express KRT5 or SFTPC.

Scale bars, 100 µm.

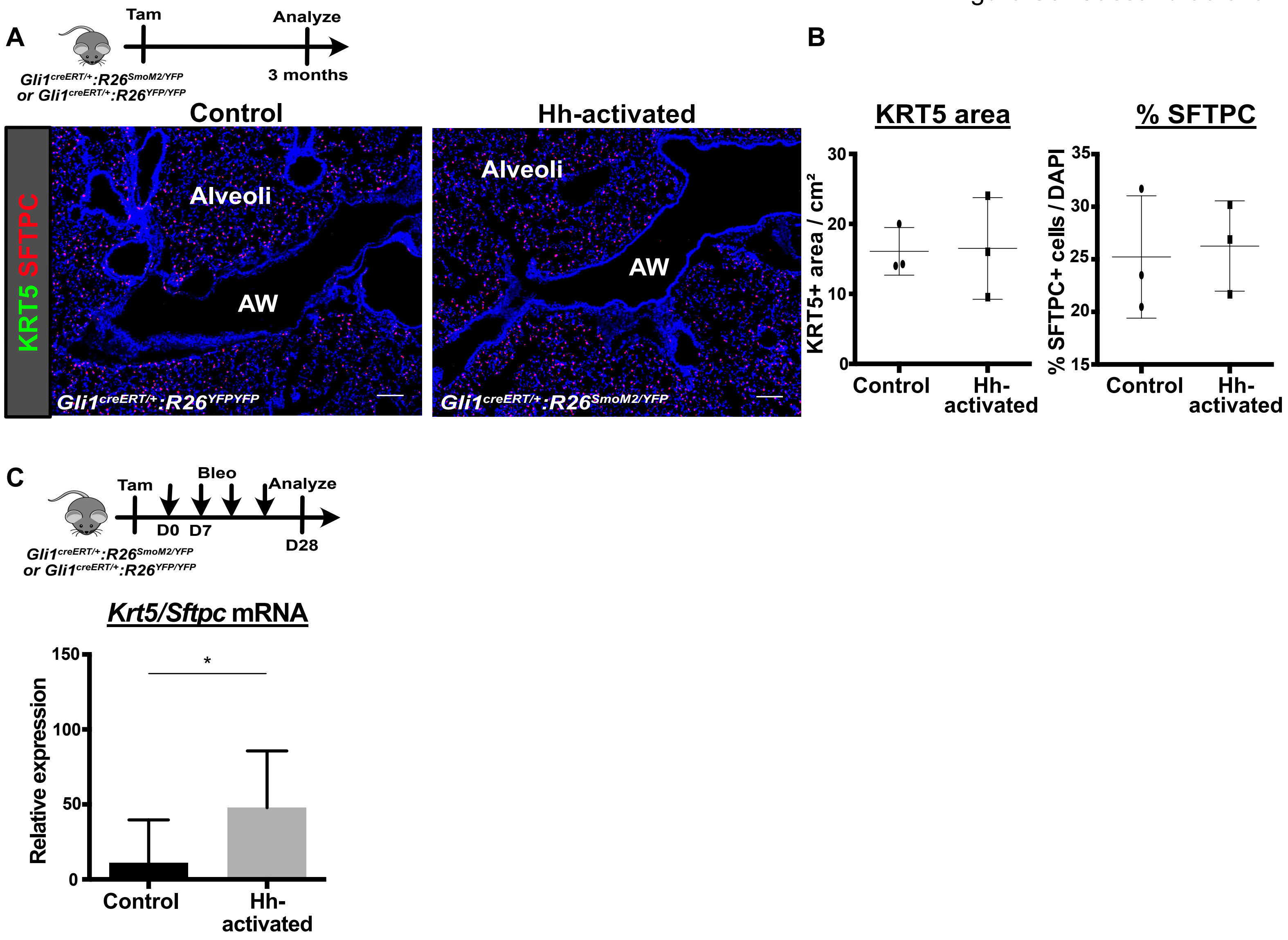
A
Tam
Analyze
3 months
Gli1creERT/+:R26SmoM2/YFP
or Gli1creERT/+:R26YFP/YFP
Control
Hh-activated
KRT5 SFTPC
Alveoli
AW
Gli1creERT/+:R26YFPYFP
Gli1creERT/+:R26SmoM2/YFP
B
KRT5 area
KRT5+ area / cm²
% SFTPC
% SFTPC+ cells / DAPI
Control
Hh-activated
C
Tam
Bleo
Analyze
D0 D7
D28
Krt5/Sftpc mRNA
Relative expression
*
Control
Hh-activated

**Figure S3. Characterization of Hh-activation during homeostasis and in fibrotic repair, related to Figure 2.**

(A) During homeostasis, histology of lungs with Hh activation in *Gli1* Lin+ cells shows same expression pattern of SFTPC and KRT5 as control lungs. No KRT5 expression is detected in the distal airways or alveoli in either condition.

(B) Histology quantification shows similar expression of total KRT5 area and percentage of SFTPC+ cells in the alveoli between Hh-activated and control lungs. Data are expressed as mean ± SD.

(C) Following fibrotic injury, qPCR analysis of the whole lung shows a significant increase in the ratio of *Krt5/Sftpc* transcripts after Hh activation of *Gli1* Lin+ cells. Data are expressed as mean ± SD.

AW = airway. Scale bars, 100 μm. See also Table S1.

A

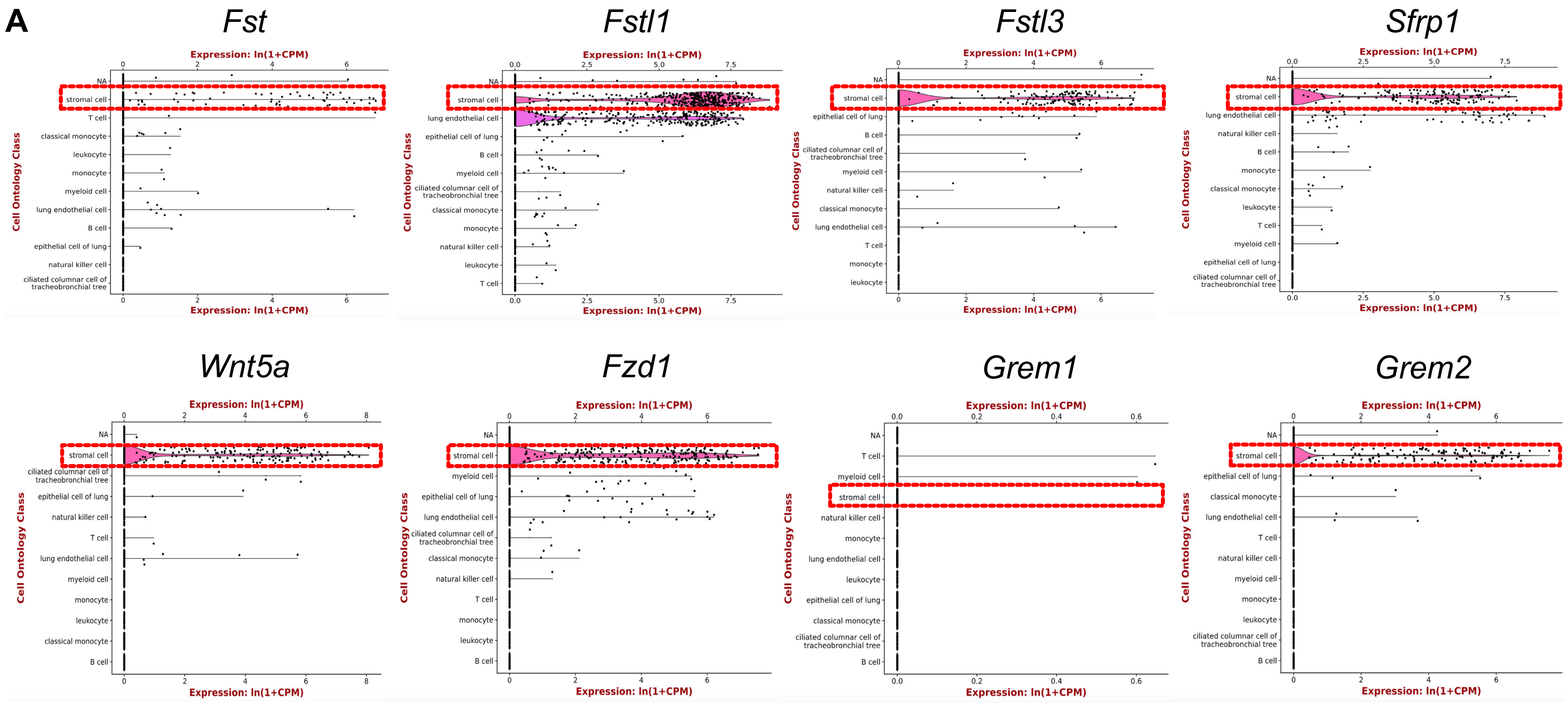
Fst
Fstl1
Fstl3
Sfrp1
Wnt5a
Fzd1
Grem1
Grem2
Expression: ln(1+CPM)
Cell Ontology Class
NA
stromal cell
T cell
classical monocyte
leukocyte
monocyte
myeloid cell
lung endothelial cell
B cell
epithelial cell of lung
natural killer cell
ciliated columnar cell of tracheobronchial tree

B

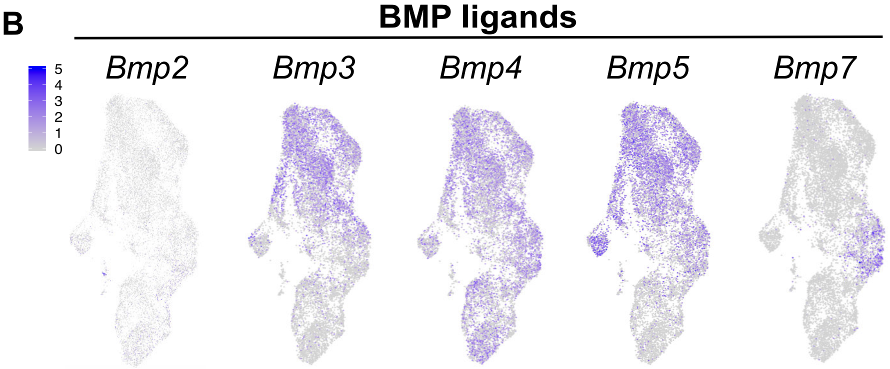
BMP ligands
Bmp2
Bmp3
Bmp4
Bmp5
Bmp7
5
4
3
2
1
0

**Figure S4. BMP antagonists and ligands are expressed in the mesenchyme, related to Figure 3.**

(A) Transcriptome analysis of whole mouse lung using publicly available cellular atlas (Tabula muris) shows secreted BMP antagonists are predominantly expressed by the mesenchyme/stroma.

(B) Expression of BMP ligands in the *Gli1* Lin+ mesenchyme.

**A**

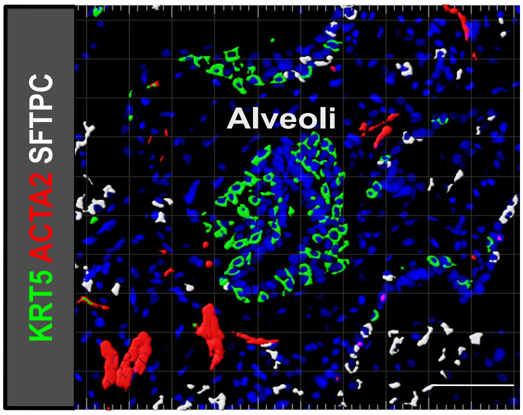
KRT5 ACTA2 SFTPC
Alveoli

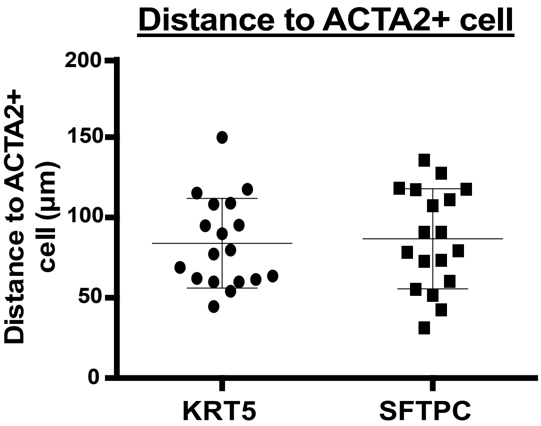
Distance to ACTA2+ cell
Distance to ACTA2+ cell (μm)
200
150
100
50
0
KRT5
SFTPC

**Figure S5. Metaplastic expression of KRT5+ basal cells and ACTA2+ fibroblastic foci in the alveoli of IPF lungs, related to Figure 6.**

(A) Ectopic expression of KRT5 and ACTA2 in the alveoli of IPF lungs form honeycomb cysts and fibroblast foci, respectively.

(B) Average cell-to-cell distance shows equivalent proximity of metaplastic KRT5 and endogenous SFTPC cells to ACTA2+ fibroblastic foci. Data are expressed as mean ± SD.

Scale bars, 100 μm.

Primers for qRT-PCR (mouse)

| mRNA Transcript | Forward Primer (5'-3') | Reverse Primer (5'-3') | Source |
|---|---|---|---|
| KRT5 | TCCAGTGTGTCCTTCCGAAGT | TGCCTCCGCCAGAACTGTA | Vaughan et al., 2015 |
| SFTPC | ATGGACATGAGTAGCAAAGAGGT | CACGATGAGAAGGCGTTTGAG | Vaughan et al., 2015 |
| FST | GATCTTGCAACTCCATCTCGGA | ACACTGAACATTGGTGGAGAGTT | NM_001301373 |
| FSTL1 | CGAGCACGATGTGGAAACGA | ACATTGGCGCAGATCTTGGA | NM_008047 |
| FSTL3 | CTGCCTCCCCTGCAAAGATTC | CGGTACATGACGCGCAAGT | Primerbank |
| GREM1 | CTGGAGACCCAGAGTACCGT | TGCGGTGCGATTCATTCTGT | NM_011824 |
| GREM2 | CTCGTCATTGCAGGATGTTCTGG | CTTGTAAGGCGAGGGGATGG | NM_011825 |
| SFRP1 | GGAAGCCTCTAAGCCCCAAG | CATCCTCAGTGCAAACTCGC | NM_013834 |
| BMP3 | GCTCTATGACAGGTACAGCGG | GTTTGAGGAGTTCCTGCGGCT | NM_001310677 |
| BMP4 | AACCAATGAGACACCATGATTCC | GAAGTGTCGCCTCGAAGTCC | NM_007554 |
| BMP5 | CTTCTTCGATCCGTGAGAGCA | GTGCTATGATCCAGTCCTGC | NM_007555 |
| BMP7 | ACGGACAGGGCTTCTCCTAC | ATGGTGGTATCGAGGGTGGAA | Primerbank |
| WNT5A | ATGCAGTACATTGGAGAAGGTG | CGTCTCTCGGCTGCCTATTT | Primerbank |
| FZD1 | CAGCAGTACAACGGCGAAC | GTCCTCCTGATTCGTGTGGC | Primerbank |
| GLI1 | CGC CCC GAC GGA GGT CTC TT | GCT GGC CGT CCC AAC TGC TT | NM_010296 |
| GAPDH | CCCCAGCAAGGACACTGAGCAAGAG | GGCCCCTCCTGTTATTATGGGGGT | Wang et al., 2018 |

**Table S1. List of qRT-PCR primers used for mouse mRNA analysis. Related to Figures 2, 4, 5, and Figure S3.**
